# Abundance and distribution of ringed and bearded seals in the Chukchi Sea: a reference for future trends

**DOI:** 10.1101/2024.08.27.608330

**Authors:** Peter L. Boveng, Vladimir I. Chernook, Erin E. Moreland, Paul B. Conn, Irina S. Trukhanova, Michael F. Cameron, Cynthia L. Christman, Justin A. Crawford, Lois Harwood, Benjamin X. Hou, Stacie M. Koslovsky, Jessica M. Lindsay, Denis I. Litovka, Josh M. London, Brett T. McClintock, Nikita Platonov, Lori Quakenbush, Erin L. Richmond, Alexander Vasiliev, Andrew L. Von Duyke, Amy Willoughby

## Abstract

Ringed (*Pusa hispida*) and bearded (*Erignathus barbatus*) seals are vulnerable to decreasing sea ice habitat in the rapidly warming Arctic. In April and May of 2016, we conducted an aerial survey over the ice-covered areas of the Chukchi Sea using thermal and color cameras to detect and count these seals on sea ice. We related the seal counts to environmental variables, and used the relationships to estimate the species’ distributions and abundance throughout the Chukchi Sea. We accounted for incomplete detection due to seals missed by sensors or image processing errors, behavioral responses to aircraft, or incomplete availability (i.e. seals that are in water or in snow dens on the ice, called lairs). For the latter, we used wet/dry records from satellite-linked bio-loggers, and remotely sensed snow melt dates to estimate the proportion of ringed seal individuals that were visible on ice for each day and location surveyed. To our knowledge, this is the first study where use of lairs by ringed seals has been formally addressed while estimating abundance from aerial surveys. Ringed seal abundance was estimated as 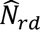 = 592,577 (95% CI: 478,448–733,929), with highest densities near Kotzebue Sound, Alaska, USA. Bearded seal abundance was estimated as 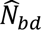 = 147,421 (95% CI: 114,155–190,380), with highest densities in broken pack ice near Bering Strait. The influence of environmental variables, such as snow depth and ice type, was consistent with prior studies of the species’ natural history, particularly ringed seals’ preference for snow of adequate depth for lairs. Our study provides the first comprehensive abundance estimates for ringed and bearded seals in the Chukchi Sea and establishes a reference for monitoring how their populations respond to Arctic warming.

## 1 Introduction

Arctic ringed seals (*Pusa hispida hispida*) and Pacific bearded seals (*Erignathus barbatus nauticus*) are denizens of the sea ice, traditional staple resources for Arctic Indigenous communities, and crucial prey for polar bears (*Ursus maritimus*). They are considered ice-associated species because they rely on sea ice to support vital aspects of their life history. Climate-driven declines in seasonal sea ice (e.g. Stroeve and Notz 2018; Stabeno and Bell 2019; Kim et al. 2023) have prompted concern for the viability of many Arctic marine mammal species (Laidre et al. 2015), including ice-associated seals. In 2012, the Arctic subspecies of ringed seals and the Beringia Distinct Population Segment (DPS) of Pacific bearded seals were listed as Threatened under the U.S. Endangered Species Act (National Marine Fisheries Service 2012a; National Marine Fisheries Service 2012b). The listing decisions were primarily based on concerns of future habitat loss (i.e. sea ice) rather than documented population declines, as trends in population status of these currently abundant species remain largely unknown (Laidre et al. 2015).

Despite concerns about ringed and bearded seal long-term persistence, it remains difficult to reliably estimate abundance and trends due to the challenges of surveys over vast and remote ranges. Different survey methods have been used over several decades, further making temporal comparisons difficult. Early aerial surveys for ringed seals in Alaska waters (e.g., the Chukchi and Beaufort seas; see Frost et al. 1988; Frost et al. 2004, and references therein) used fixed-wing aircraft at low altitude, with human observers visually counting and identifying seals in predetermined survey strips over the landfast ice zone. A shared set of survey protocols allowed these surveys to be compared to one another as indices of ringed seal abundance. However, variation in factors that determine a seal’s detection probability make it difficult to compare those early surveys with more recent surveys. These factors include aircraft disturbance of seals, and, especially, availability and perceptibility (Marsh and Sinclair 1989). For aerial surveys of Arctic seals, availability is the probability that a seal is ‘hauled out’ of the water *and* exposed on the ice or snow surface for perception by airborne observers or sensors. Perceptibility is the probability of a seal being perceived by an observer or sensing process, given that the seal is available.

For aerial surveys of ringed and bearded seals in 1999 and 2000, Bengtson et al. (2005) used distance sampling methods (Buckland et al. 2001) to count seals, and wet/dry (‘haul-out’) records from satellite-linked bio-loggers to estimate and correct for the proportion of ringed seals that were not available because they were in the water; there was no correction for ringed seals hidden under snow, and there were no haul-out records for bearded seals. The ringed seal abundance estimates, partially corrected for availability, are likely more robust for comparison purposes but, like the earlier surveys, they remained uncorrected for perceptibility on the track line (g(0) in distance sampling terminology; e.g., Conn et al. 2012; Burt et al. 2014). In the mid-2000s, helicopter-based surveys of ice-associated seals in the Bering Sea attempted to account for g(0) < 1.0 by using mark-recapture distance sampling (Borchers et al. 2006) to estimate perceptibility, and haul-out records to estimate availability (Conn et al. 2013; Ver Hoef et al. 2014). However, due to seals’ response to noise, helicopter surveys caused disturbance that negatively biased abundance estimates (M. Cameron, pers. obs.) and contributed to substantial species identification errors (Conn et al. 2013).

In 2012 and 2013, our team of U.S. and Russian Federation scientists conducted the first comprehensive surveys of ringed, bearded, spotted (*Phoca largha*), and ribbon (*Histriophoca fasciata*) seals in the Bering Sea and Sea of Okhotsk (Chernook et al. 2014; Conn et al. 2014; Sigler et al. 2015). These sensor-based aerial surveys used fixed-wing aircraft, but instead of human observers they used thermal imaging to detect the warm seals resting on the cold sea ice. Thermal detections were then matched with higher-resolution digital color photographs to determine species. As far as we are aware, this was the first effort to estimate absolute seal abundance while accounting simultaneously for incomplete perception by the sensor process, species misidentification, and availability less than 1.0. However, ringed seal abundance could not be corrected for availability (Conn et al. 2014) because of uncertainty about the proportion of ringed seals that were using snow dens on the sea ice (‘subnivean lairs’; Kelly et al. 2006) while surveys were conducted.

In 2016, our team conducted a second joint U.S.–Russian Federation survey, for ringed and bearded seals in the Chukchi Sea and a small portion of the East Siberian Sea, again using sensor-based protocols in fixed-wing aircraft. That survey is the focus of this paper, in which we report results of the Chukchi and East Siberian Surveys (ChESS), using a spatio-temporal statistical model to estimate the abundance of ringed and bearded seals while accounting for availability, perceptibility, and behavioral response of seals to survey aircraft. Our approach incorporates a novel conceptual model for seal availability that relies on haul-out data for both ringed and bearded seals, and on the timing of snow melt onset to help quantify ringed seals’ transition from hauling out in lairs to hauling out on the exposed ice or snow surface—an important component of their availability.

## 2 Methods and Materials

### 2.1 Overview

In our aerial survey, we obtained counts of ringed and bearded seals in the area sampled by our sensors. With the survey area divided into grid cells, seal counts within grid cells were then related to habitat variables (covariates) and used to predict abundances in unsampled locations. Our approach relied on maximum likelihood, and took into account several nuisance processes that affected the detectability of seals: incomplete availability of seals in the water or under snow, incomplete perception of seals by the sensors or image processing method, and behavioral response of seals to aircraft (i.e. escape behavior). Our main quantities of interest were the values of ringed and bearded seal abundance that maximized this likelihood, given the observed counts, but the covariate relationships are also informative and help to describe habitat use. In the remainder of this section, we describe the survey and image processing methods, our choices of explanatory variables, and our approaches to modeling ringed and bearded seal density and availability.

### 2.2 Survey methods

Two teams of researchers, one in the Russian Federation exclusive economic zone (EEZ) and one in the U.S. EEZ, conducted visual and instrument-based aerial surveys over the Chukchi Sea during April and May of 2016 (Figure 1). Surveys in the Russian EEZ (Chernook et al. 2019) were conducted from an AN-26 Arctica aircraft equipped with a Malakhit-M cooled long-wavelength infrared (LWIR; 8–14 μm) scanner; cooled infrared sensors provide greater sensitivity than uncooled sensors. At a target altitude of 250 m, the LWIR scanner had a ‘ground sampling distance’ (GSD; i.e. resolution) of 11 cm/pixel on the center of the continuously recorded image strip. Three Nikon D800 digital color 36-megapixel cameras with 50-mm lenses were mounted at angles such that their combined fields of view completely covered the strip width of the LWIR scanner, which was 483 m when flying at a target altitude of 250 m. The center-mounted color camera had a GSD of 2.27 cm/pixel at the center of its image.

**Figure 1.**
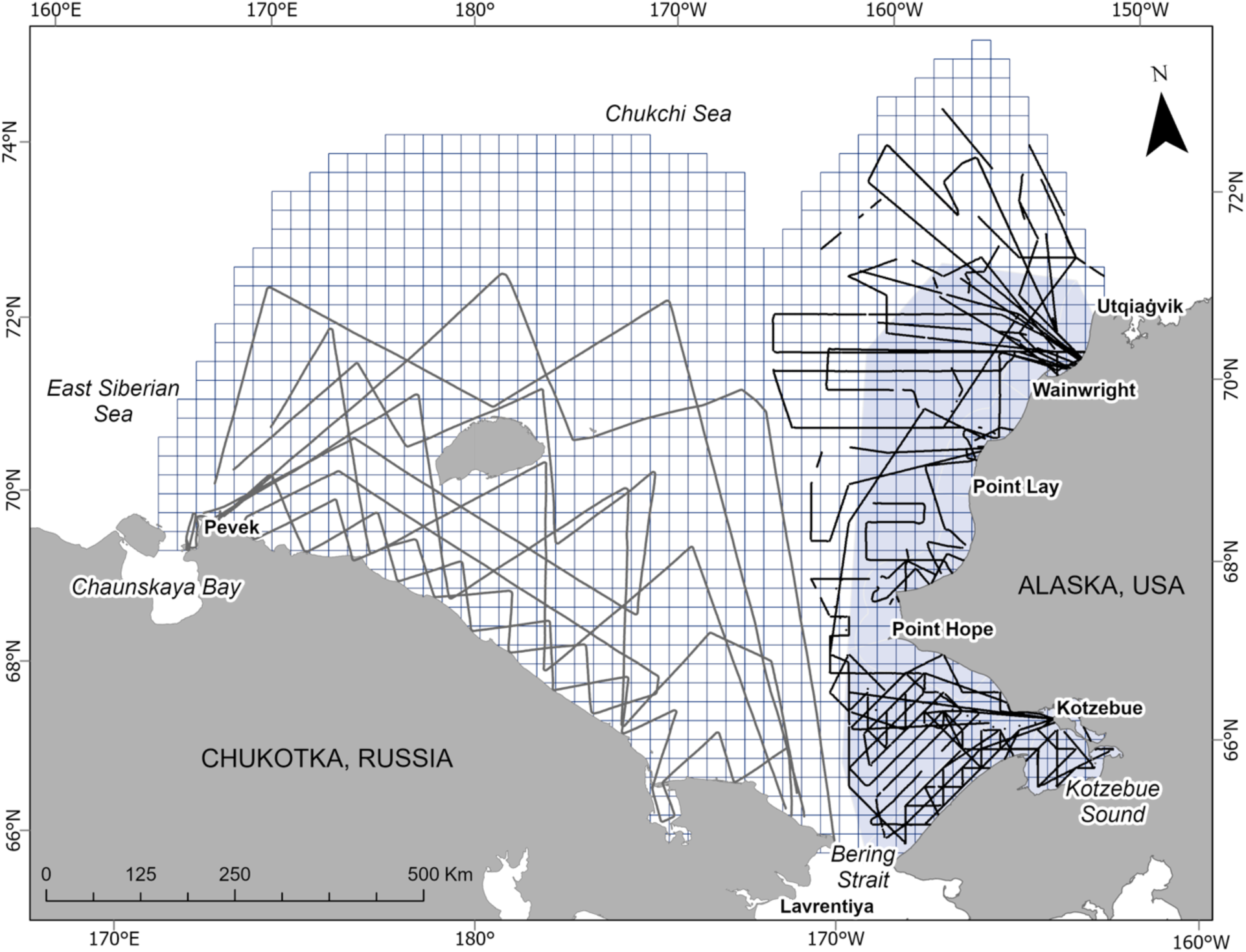
The ringed and bearded seal aerial survey area in the Chukchi Sea, 2016. The study area is divided into 25 × 25 km (625 km^2^) grid cells (blue lines). Black lines represent ‘on effort’ flight tracks over U.S. waters and dark gray lines represent flight tracks over Russian Federation waters. Gaps in lines represent ‘off effort’ segments, primarily due to fog. Blue shading represents the 1999 and 2000 seal survey area (Bengtson et al. 2005).

Surveys in the U.S. EEZ were conducted from a King Air A90 aircraft configured with three downward facing, cooled LWIR cameras (FLIR A6751sc SLS) fitted with 25-mm lenses mounted in the open-air belly port. One uncooled LWIR camera (FLIR A645sc) was used in place of a cooled camera for 13 flights while the cooled camera was temporarily inoperable. Both types of LWIR camera provided 20–23 cm GSD at the target survey altitude of 305 m. We paired each LWIR (hereafter referred to as ‘thermal’) camera with a 29-mega-pixel machine-vision color camera (Prosilica GT6600c) fitted with a 100-mm Zeiss lens to provide a GSD of 1.71–2.13 cm/pixel at survey altitude. Paired color and thermal images were collected continuously while on survey effort at a rate of two frames per second. Dedicated on-board computers running custom software (Skeyes; MoviTHERM, Irvine, CA) managed image acquisition and synchronized data logging from an integrated Arduino GPS. At the target altitude, the thermal cameras covered a strip width of 470 m. The color images, which determined the area surveyed, had a footprint width equal to 83% of the thermal strip width, or 390 m. The target survey ground speed of approximately 260km/h was set to maximize the range of the aircraft, and resulted in 50% forward overlap of images. Side camera pairs were set at an internal angle of 25.5° to allow a slight gap between sea ice coverage in the images collected from the center and side cameras, maximizing survey swath while eliminating image processing and analysis complications from duplicate detections.

Variable weather and sea ice conditions make it difficult to implement a pre-defined sampling design. Therefore, the survey teams were instructed to spread survey effort over space and time, as is commonly done in space-filling designs developed for spatial modelling (Diggle and Lophaven 2006). This model-based approach exhibited satisfactory performance in initial simulation testing, and higher performance with survey effort distributed over space and time, compared to stratified, design-based approaches used in previous surveys (Conn et al. 2016). Survey teams were instructed to avoid consideration of expected seal density or habitat quality when laying out or modifying survey track lines, to minimize bias due to preferential sampling (Diggle et al. 2010; Conn et al. 2017).

We divided our study area into 1,354 grid cells of approximately 25 × 25 km (625 km^2^) each (Figure 1), commensurate with the resolution of sea ice concentration data that we related to seal densities (see **2.4 Explanatory variables**). We tallied seal counts and survey effort at this scale, though with slightly different methods for U.S. and Russian survey flights. In the U.S. surveys, we quantified effort as the total area photo-graphed with the color cameras, and did not attempt to model thermal detections that fell outside the color images. For the Russian surveys, we used the area of the thermal scanner swath to quantify effort, because photographs were only triggered when thermal signatures characteristic of seals on ice (‘hot spots’) were detected on the thermal scanner; not all hot spots were photographed, which led to different probabilities of identifying detected seals to species, compared to the U.S. surveys. Methods to account for these differences are described in the section on abundance modelling below. The seal counts and metadata we used from the Russian surveys were published in Chernook et al. (2019). Our analysis combines the U.S. and Russian data under a single, more comprehensive model for estimation of seal abundance and distribution across the Chukchi Sea.

### 2.3 Processing images for seal detection

#### 2.3.1 Russian Federation surveys

The thermal scanner recording was processed with custom software to automatically mark seals on the ice using a combination of temperature contrast, size, and shape of hot spots. Operators adjusted software settings during the process to ensure the reliable identification of thermal signatures from seals while excluding false positives, for example caused by the water in seal access holes or dirt (Chernook et al. 2000a; Chernook et al. 2000b). Operators reviewed and manually corrected the results of automatic thermal processing and used detection times and coordinates to match thermal detections with the corresponding color images. On color images, seals detected as hot spots from the thermal scanner were marked by operators, and remaining false-positive hot spots were eliminated by verification with the content of the color images. Combined thermal and color image processing was independently conducted by two counters. Results from both counts were compared for discrepancies and resolved. Once a seal was found in both thermal and color images, a digital portrait was provided to biologists for species identification and determination of age class (pup or non-pup). This processing resulted in a database containing unique seal identifiers, species information, geographical coordinates, detection times, and thermal and color image file names.

#### 2.3.2 U.S. surveys

After surveys were completed, we used a partially automated process to search for and classify seals in the survey images. In the first step, the same software used for image acquisition was used to apply an automated algorithm to find hot spots in the thermal images. The algorithm was designed to exclude hot spots that were either too small in area (< 6 pixels) or too large (>300 pixels), and to use simple measures of the hot spot temperature profile to further eliminate false positive detections. Specifically, a curve was fit to the grayscale pixel values along a single line bisecting the warmest region of each hot spot. The peak value, the maximum slope along the curve, and the second derivative of the curve were compared to threshold values that had been empirically derived from thermal signatures of known seals in aerial imagery collected prior to our survey. This algorithm identified 316,099 candidate hot spots from 997,642 processed thermal images. In the second step, all candidate hot spots were then reviewed visually to remove those that were unlikely to be from seals, reducing the number that required further processing. For each of the remaining 15,454 hot spots, the corresponding color image was examined by experienced observers to determine the hot spot’s source and to classify species and age class (pup or non-pup) of detected seals. To quantify the seal perception rate of the thermal sensors and this image process, a systematic random sample of 28,052 color images was visually searched for the presence of seals by an observer without access to the thermal images. The partially automated process detected 247 of the 258 seals found in the visual search, for an overall perception probability of 0.96 (Supplement 1). Lacking a similar study of the sensors and perception process used in the Russian Federation surveys, we applied the same perception probability to the Russian data.

### 2.4 Explanatory variables

Counts of seals present on sea ice are a function of seal density and the probability of detection, which has components of availability and perceptibility (Marsh and Sinclair 1989). We selected 14 environmental, temporal, and geographic covariates to help explain variation in density and availability (Table 1). Based on Indigenous knowledge (e.g., Oceana and Kawerak Inc. 2014; Huntington et al. 2015), natural history observations (e.g., Burns 1981), previous aerial surveys (e.g., Bengtson et al. 2005), and satellite tracking results (e.g., Crawford et al. 2012; Olnes et al. 2020; Von Duyke et al. 2020), we anticipated that seal densities would vary by sea ice concentration and type, with ringed seals typically selecting landfast and nearshore pack ice, and bearded seals typically selecting pack ice. We also anticipated that both species would occur at higher densities south of Point Hope (Bengtson et al. 2005; Von Duyke et al. 2020). Ringed seals require sufficient snow to construct subnivean lairs, so we included a remotely sensed snow depth covariate. We included wind speed, air temperature, precipitation, atmospheric pressure, day of year, and solar hour as availability covariates because more ringed and bearded seals are known to haul out on ice in calm, warm, dry conditions, later in the spring, and at hours closer to solar noon and midnight (Burns and Harbo 1972; Frost et al. 2004; London et al. 2024).

**Table 1.**
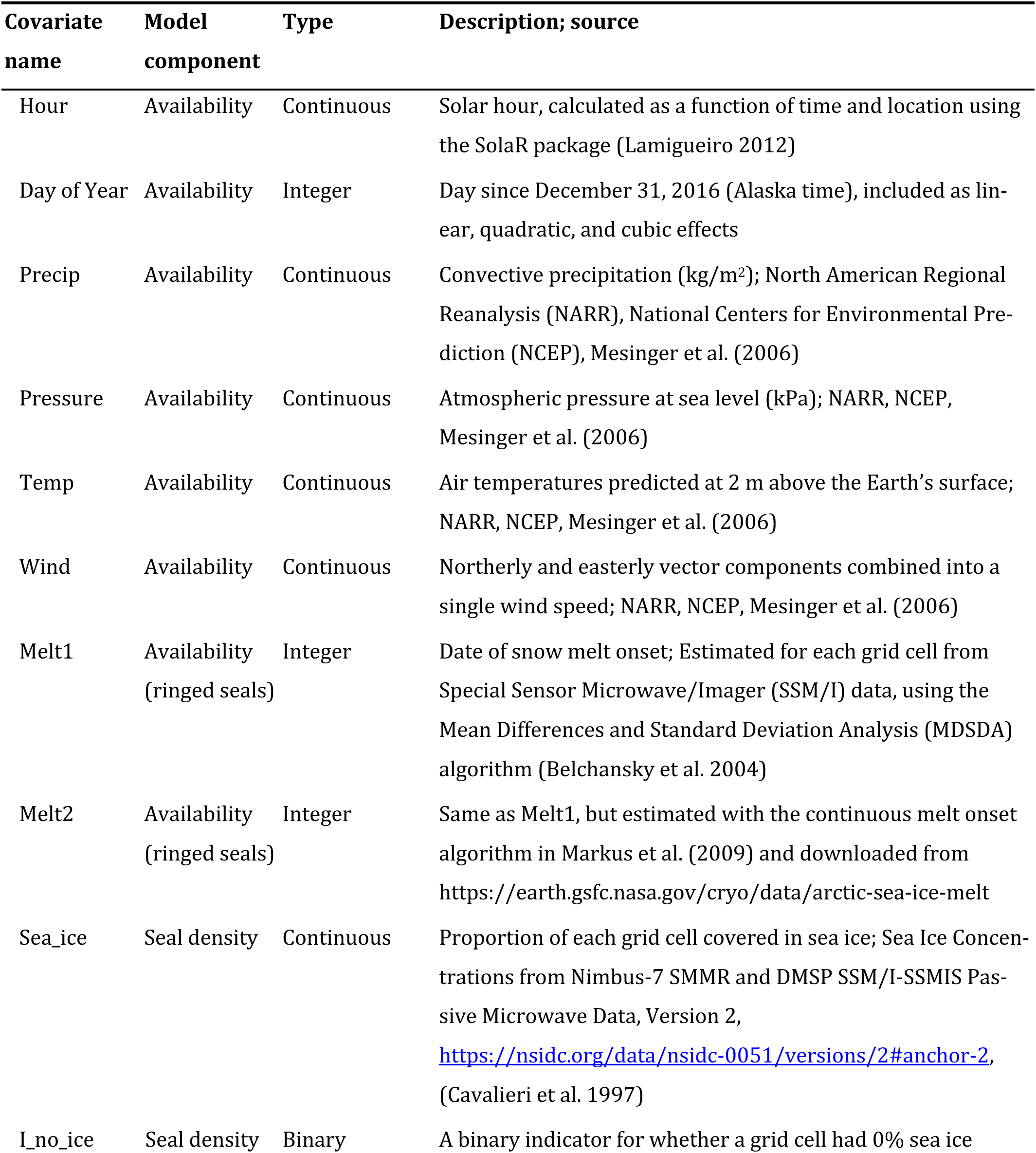
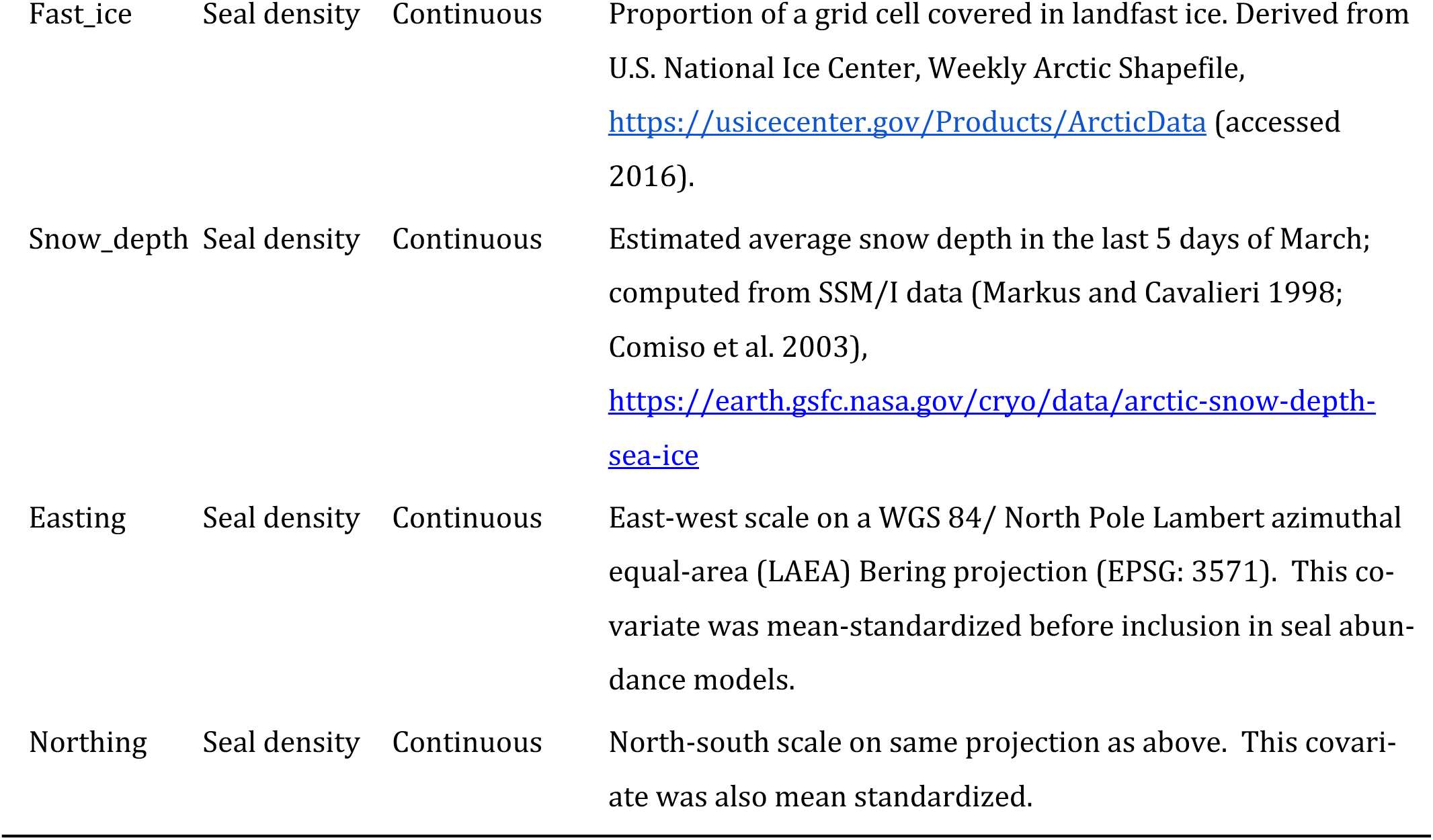
Explanatory covariates used in the analysis of ringed and bearded seal availability and density (abundance).

| **Covariate name** | **Model component** | **Type** | **Description; source** |
| --- | --- | --- | --- |
| Hour | Availability | Continuous | Solar hour, calculated as a function of time and location using the SolaR package (Lamigueiro 2012) |
| Day of Year | Availability | Integer | Day since December 31, 2016 (Alaska time), included as linear, quadratic, and cubic effects |
| Precip | Availability | Continuous | Convective precipitation (kg/m<sup>2</sup>); North American Regional Reanalysis (NARR), National Centers for Environmental Prediction (NCEP), Mesinger et al. (2006) |
| Pressure | Availability | Continuous | Atmospheric pressure at sea level (kPa); NARR, NCEP, Mesinger et al. (2006) |
| Temp | Availability | Continuous | Air temperatures predicted at 2 m above the Earth's surface; NARR, NCEP, Mesinger et al. (2006) |
| Wind | Availability | Continuous | Northerly and easterly vector components combined into a single wind speed; NARR, NCEP, Mesinger et al. (2006) |
| Melt1 | Availability<br>(ringed seals) | Integer | Date of snow melt onset; Estimated for each grid cell from Special Sensor Microwave/Imager (SSM/I) data, using the Mean Differences and Standard Deviation Analysis (MDSDA) algorithm (Belchansky et al. 2004) |
| Melt2 | Availability<br>(ringed seals) | Integer | Same as Melt1, but estimated with the continuous melt onset algorithm in Markus et al. (2009) and downloaded from https://earth.gsfc.nasa.gov/cryo/data/arctic-sea-ice-melt |
| Sea_ice | Seal density | Continuous | Proportion of each grid cell covered in sea ice; Sea Ice Concentrations from Nimbus-7 SMMR and DMSP SSM/I-SSMIS Passive Microwave Data, Version 2, https://nsidc.org/data/nsidc-0051/versions/2#anchor-2, (Cavalieri et al. 1997) |
| I_no_ice | Seal density | Binary | A binary indicator for whether a grid cell had 0% sea ice |
| Fast_ice | Seal density | Continuous | Proportion of a grid cell covered in landfast ice. Derived from U.S. National Ice Center, Weekly Arctic Shapefile, https://usicecenter.gov/Products/ArcticData (accessed 2016). |
| Snow_depth | Seal density | Continuous | Estimated average snow depth in the last 5 days of March; computed from SSM/I data (Markus and Cavalieri 1998; Comiso et al. 2003), https://earth.gsfc.nasa.gov/cryo/data/arctic-snow-depth-sea-ice |
| Easting | Seal density | Continuous | East-west scale on a WGS 84/ North Pole Lambert azimuthal equal-area (LAEA) Bering projection (EPSG: 3571). This covariate was mean-standardized before inclusion in seal abundance models. |
| Northing | Seal density | Continuous | North-south scale on same projection as above. This covariate was also mean standardized. |

Accounting for ringed seals’ availability is further complicated by their seasonal use of lairs, wherein they are not visible to the aerial survey sensors. Their transition from hauling out in lairs to hauling out on the exposed surface of the ice or snow (i.e. basking) is often not abrupt. This transition is partly influenced by increasing ambient temperatures that support the seals’ molt (Feltz and Fay 1966) and cause snow melt, eventually leading to lair collapse (Kelly et al. 2006). Individuals may haul out on the ice to bask near breathing holes prior to lair collapse, and may return to using lairs between basking bouts. Ringed seals will also dig out before lairs collapse when conditions (i.e. air temperature, wind speed, and solar energy) are favorable for basking, but some lairs remain in use late into spring (Kelly and Quakenbush 1990). Thus, a seal’s transition away from hauling out in lairs can be a gradual process—under normal conditions—when lairs remain intact, or more abrupt when rain, thin snow, or early snow melt cause lairs to collapse prematurely. We refer to this transition as emergence (Lindsay et al. 2021). To explore a relationship between emergence and snow melt we tested two remotely sensed covariates (Table 1) associated with snow melt onset, the date on which the snow first becomes saturated with moisture.

### 2.5 Availability modeling

Our surveys only had the ability to detect seals that were basking on ice and visible to the airborne sensors. Thus, to estimate seal abundance from aerial survey counts, we needed to account for the proportion of seals that were available, *a_ist_*. Here, the subscripts indicate that availability has the potential to vary by species (*i*), location (i.e. grid cell; *s*), and time (*t*). An alphabetical list of notation used in model development is provided in Table 2.

**Table 2.**
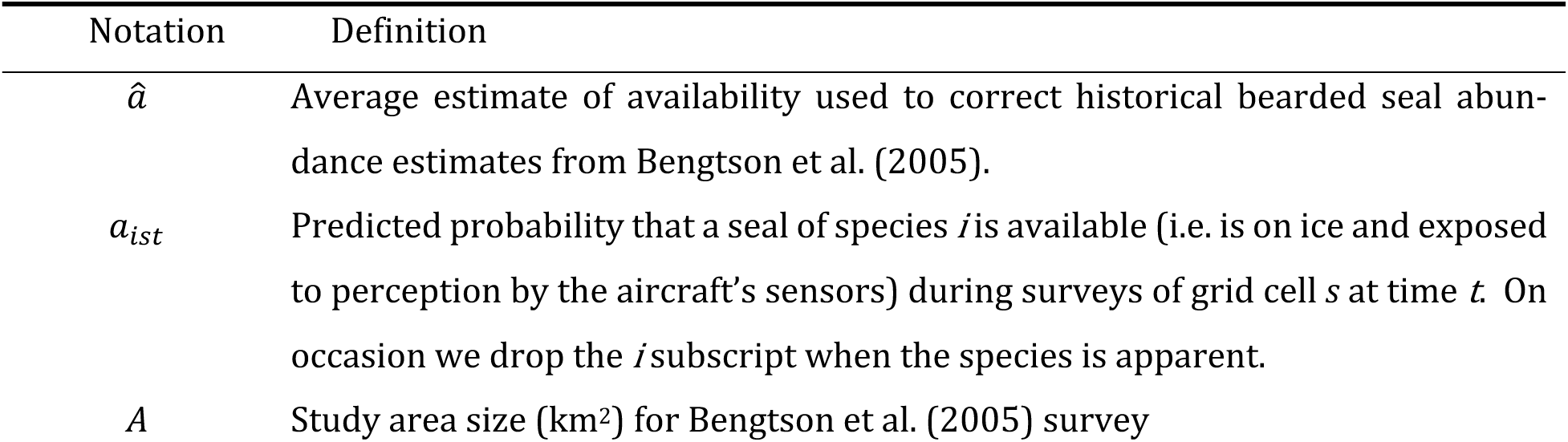
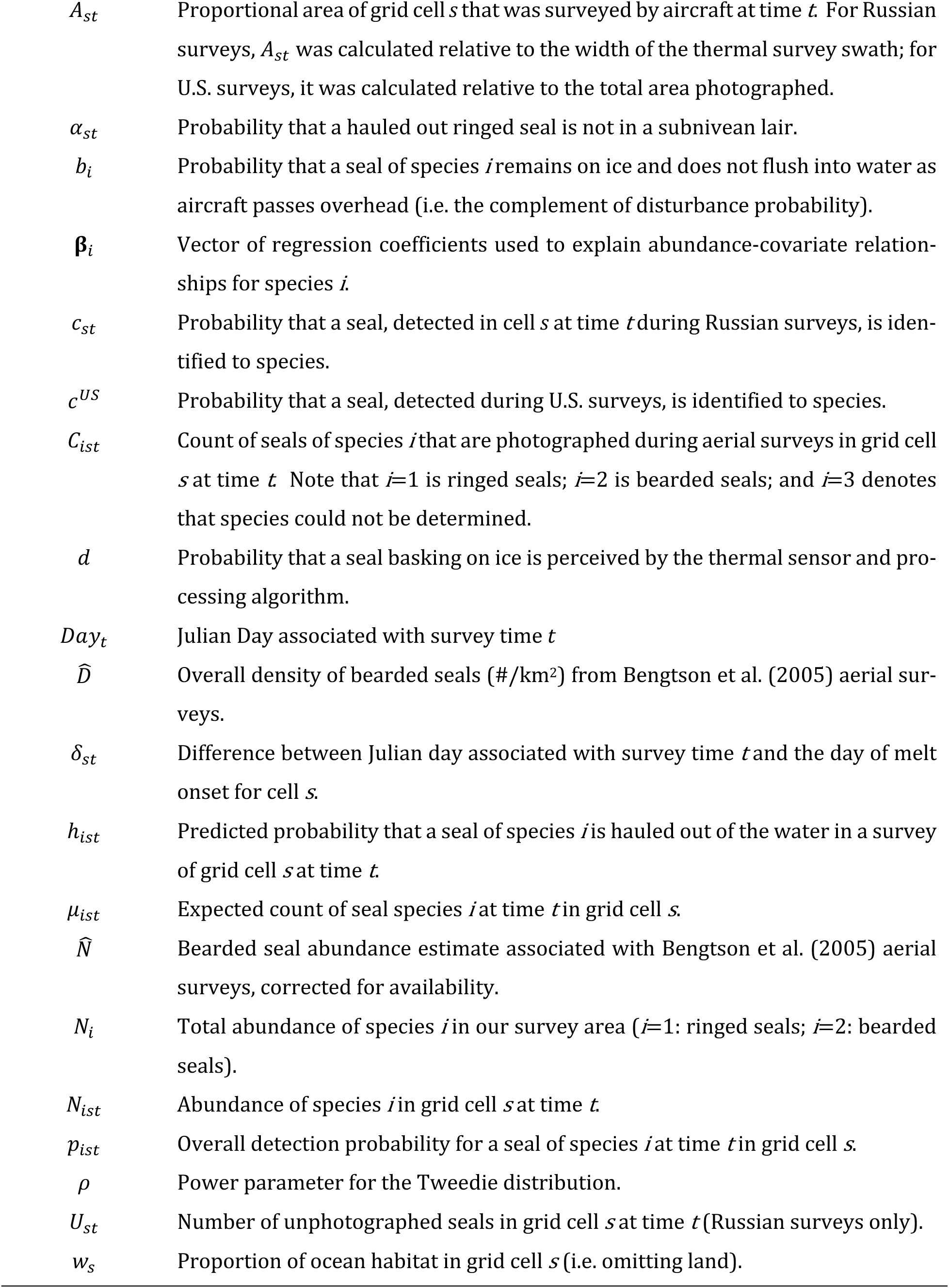
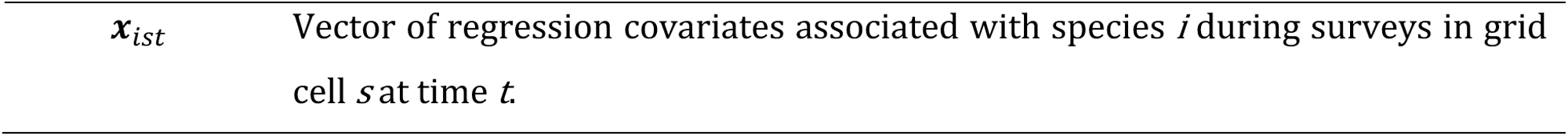
Notation for models fitted to ringed and bearded seal aerial survey count data.

| Notation | Definition |
| --- | --- |
| $\hat{a}$ | Average estimate of availability used to correct historical bearded seal abundance estimates from Bengtson et al. (2005). |
| $a_{ist}$ | Predicted probability that a seal of species $i$ is available (i.e. is on ice and exposed to perception by the aircraft's sensors) during surveys of grid cell $s$ at time $t$. On occasion we drop the $i$ subscript when the species is apparent. |
| $A$ | Study area size (km<sup>2</sup>) for Bengtson et al. (2005) survey |
| $A_{st}$ | Proportional area of grid cell $s$ that was surveyed by aircraft at time $t$. For Russian surveys, $A_{st}$ was calculated relative to the width of the thermal survey swath; for U.S. surveys, it was calculated relative to the total area photographed. |
| $\alpha_{st}$ | Probability that a hauled out ringed seal is not in a subnivean lair. |
| $b_i$ | Probability that a seal of species $i$ remains on ice and does not flush into water as aircraft passes overhead (i.e. the complement of disturbance probability). |
| $\beta_i$ | Vector of regression coefficients used to explain abundance-covariate relationships for species $i$. |
| $c_{st}$ | Probability that a seal, detected in cell $s$ at time $t$ during Russian surveys, is identified to species. |
| $c^{US}$ | Probability that a seal, detected during U.S. surveys, is identified to species. |
| $C_{ist}$ | Count of seals of species $i$ that are photographed during aerial surveys in grid cell $s$ at time $t$. Note that $i=1$ is ringed seals; $i=2$ is bearded seals; and $i=3$ denotes that species could not be determined. |
| $d$ | Probability that a seal basking on ice is perceived by the thermal sensor and processing algorithm. |
| $Day_t$ | Julian Day associated with survey time $t$. |
| $\hat{D}$ | Overall density of bearded seals ($\#/km^2$) from Bengtson et al. (2005) aerial surveys. |
| $\delta_{st}$ | Difference between Julian day associated with survey time $t$ and the day of melt onset for cell $s$. |
| $h_{ist}$ | Predicted probability that a seal of species $i$ is hauled out of the water in a survey of grid cell $s$ at time $t$. |
| $\mu_{ist}$ | Expected count of seal species $i$ at time $t$ in grid cell $s$. |
| $\hat{N}$ | Bearded seal abundance estimate associated with Bengtson et al. (2005) aerial surveys, corrected for availability. |
| $N_i$ | Total abundance of species $i$ in our survey area ($i=1$: ringed seals; $i=2$: bearded seals). |
| $N_{ist}$ | Abundance of species $i$ in grid cell $s$ at time $t$. |
| $p_{ist}$ | Overall detection probability for a seal of species $i$ at time $t$ in grid cell $s$. |
| $\rho$ | Power parameter for the Tweedie distribution. |
| $U_{st}$ | Number of unphotographed seals in grid cell $s$ at time $t$ (Russian surveys only). |
| $w_s$ | Proportion of ocean habitat in grid cell $s$ (i.e. omitting land). |
| $\mathbf{x}_{ist}$ | Vector of regression covariates associated with species $i$ during surveys in grid cell $s$ at time $t$. |

#### 2.5.1 Ringed seal availability

A ringed seal that is hauled out on the ice but concealed within a lair is not available to be detected in aerial surveys. Therefore, we treated ringed seal availability as a product of two components: (1) the probability that a seal is out of the water (*h_st_*) and (2) the probability that it is visible on the snow or ice surface (i.e. not in a lair), given that it is out of the water (*α_st_*), such that *a_st_* = ℎ*_st_α_st_*.

To estimate ℎ*_st_* for ringed seals we used hourly wet/dry data obtained during 2005–2019 from satellite-linked bio-loggers on 65 individuals in the Bering, Chukchi, and Beaufort seas (Harwood et al. 2007; Kelly et al. 2010a; Crawford et al. 2012; Crawford et al. 2019; Von Duyke et al. 2020). Using only the data from April– June, we fit generalized additive models (GAMs) in the mgcv package (Wood 2017) to the proportion of ringed seals hauled out during aerial survey hours (09:00–15:00 local solar time) in relation to day-of-year, spring air temperature, days from snow melt onset, and sea (Bering, Chukchi, or Beaufort). For additional details on how ringed seal haul-out probabilities (ℎ*_st_*) were modeled, see Supplement section 2.

We used temporal trends in seal counts (Lindsay et al. 2021) to estimate the shape of *α_st_*. Kelly et al. (2006) showed that the proportion of radio-tagged ringed seals visible on the ice-snow surface increases as the snow becomes saturated with water, likely because lairs are melting and becoming unusable, or unneeded due to warmer ambient temperatures that promote basking. Therefore, we used snow melt onset (i.e. saturation) dates estimated with the MDSDA algorithm (Mean Differences and Standard Deviation Analysis of passive microwave brightness temperatures; Belchansky et al. 2004) as an indicator of emergence from lairs. Defining

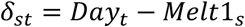

to be the difference between the survey day (*Day_t_*) and the day of snow melt onset (*Melt*1_%_) indicated by the MDSDA algorithm for cell s (such that negative values indicate days prior to snow melt onset and positive values indicate days after snow melt onset), we used this finding to set up a logistic model for the proportion of hauled-out ringed seals that have emerged from lairs:

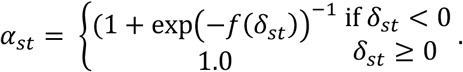

We included this emergence model in our larger model for ringed seal counts, using a penalized spline basis for *f*(*δ_st_*) to estimate its (approximately sigmoid) shape, and an additional likelihood penalty to force *α_st_* to approach 1.0 by the time the MDSDA algorithm indicated that the snow melt onset had occurred. For additional information on this conceptual model, see Supplement section 2.

#### 2.5.2 Bearded seal availability

London et al. (2024) estimated availability of bearded, ribbon, and spotted seals in the Bering and Chukchi seas. Specifically, they applied generalized linear mixed pseudo-models (Ver Hoef et al. 2010) to wet/dry records obtained in the same manner as we described for ringed seals. Because bearded seals rest on the ice surface and do not obscure themselves in snow lairs—they are available to be detected if they are hauled out during surveys. We used London et al.’s fitted model directly to predict bearded seal availability for each grid cell and time that we surveyed. Predictions depended on day-of-year, hour-of-day, temperature, wind speed, precipitation, and barometric pressure (Table 1). London et al.’s model also used latitude to describe changes in haul-out behavior from the Bering Sea, north through the southern Chukchi Sea, though because there were few bearded seal telemetry records from the middle and northern Chukchi Sea during April and May, we used a single latitude value consistent with the southern Chukchi Sea for all predictions. We also used estimated variances of predictions to propagate uncertainty about availability into standard errors of abundance estimates (see Supplement section 1).

### 2.6 Abundance (density) modelling

To estimate seal abundance, we fit a statistical model that expressed the expected count of each species as a function of (a) true, underlying abundance, (b) area surveyed, (c) the expected proportion of seals that were basking (as a function of environmental conditions), and (d) perception probability for seals that were basking on ice. We also evaluated species misclassification probabilities (Conn et al. 2013; McClintock et al. 2015), but we found that they were negligible and did not include them in the model. Our model is spatially and temporally explicit, allowing expected abundance, and thus expected counts, to change by grid cell and by day as ice and other conditions changed. We write the overall model hierarchically so that each component can be examined in sequence.

#### 2.6.1 Seal densities over space and time

To model how seal densities vary over time and space, we let *N*_i_give the total abundance in the survey area of species *i*. We follow the suggestion of Conn et al. (2015) to stabilize estimation by assuming that abundance in the survey area remains constant over the course of the survey, though we allow spatial distributions to shift within the survey area as sea ice changes. Given total abundance, we write the abundance of species *i*, in spatial location (grid cell) *s*, at time (day) *t*, as

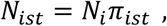

where *π_ist_* is the proportion of species *i* that is in grid cell *s* at time *t*. We then write *π_ist_* as a function of habitat and landscape covariates using a multinomial logit link function to constrain 0 ≤ *π_ist_* ≤ 1 and

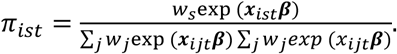

Here, *w*_s_ is the proportion of grid cell *s* that is composed of marine habitat (i.e. omitting land), ***x****_ist_* is a row vector of explanatory variables for species *i* in cell *s* at time *t*, and ***β*** is a column vector of regression coefficients. We used a parameterization that allowed for smooth effects of continuous covariates by imbedding penalized splines into estimation. This was done with the mgcv package (Wood 2017) to formulate ***x****_ist_* in terms of cubic spline bases and then penalizing ***β*** during estimation; for more detail, see Supplement section 1. We used sea ice concentration, proportion of landfast ice, easting, northing, and snow depth as covariates for seal densities. Because it is difficult to determine the effective number of parameters for this class of models, we did not conduct a formal model selection process. We provide further justification for the set of covariate predictors in **4. Discussion**.

#### 2.6.2 Expected seal counts related to abundance

Given a model for how abundance varies over space, we next describe how abundance relates to seal counts. We start by noting that (1) transects only sample a small fraction of a grid cell, (2) not all seals are detected, and (3) not all seal observations are identified to species. We thus write the expected count of each observation type (*i*=1: ringed seal; *i*=2: bearded seal; *i*=3: unknown seal species) as

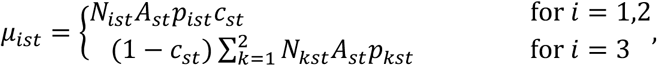

where *A_st_* is the proportion of ocean habitat in grid cell *s* that is surveyed at time *t*, *p_ist_* is the probability of detecting a seal of species *i* that is at location *s* and time *t*, and *c_st_* is the probability of identifying the seal to species. We further decompose *p_ist_* as

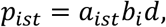

where *a_ist_* is the probability that a seal is available during the survey, *b_i_* is a non-disturbance probability (i.e. the probability that a basking seal remains on ice and is not flushed into the water before the sensors pass overhead), and *d* is the probability that a basking seal is perceived by thermal sensors and semi-automated image processing steps (Supplement section 1). We developed an informed prior distribution for *p_ist_* based on auxiliary data sources. In particular, we used disturbance trials to provide information on *b*_$_and double sampling to provide information on *d* (Conn et al. 2014).

#### 2.6.4 Classification of counts to species

Count data from the aerial surveys consisted of counts of ringed seals (*C*_1*st*_), counts of bearded seals (*C*_2*st*_), and counts of seals of unknown species (*U_st_*). The probability of identifying seals to species (*c_st_*) differed dramatically between U.S. and Russian surveys. In the U.S. surveys, we based effort on area photographed, and only 10 out of 5,720 seal observations (0.17%) were classified as ‘unknown’ species. We used a single value for *c_st_*, *c*^US^, where *c*^US^ = ∑*_s_* ∑*_t_* ∑*_i_ C_ist_*⁄∑*_s_* ∑*_t_* ∑*_i_*(*C_ist_* + *U_st_*) (here, the summation over *s* only includes cells surveyed in U.S. transects). In Russian surveys, a variable proportion of seals had clear photographs, such that 148 out of 318 seal observations (46.5%) had species information. We thus used different approaches for parameterizing *c_st_*. We set *c_st_* = ∑*_i_ C_ist_*⁄(∑*_i_ C_ist_* + *U_st_*) for cases where (∑*_i_ C_ist_* + *U_st_*) > 0. For times and cells where no seals were counted, we set *c_st_* = ∑*_s_* ∑*_t_* ∑*_i_ C_ist_*⁄∑*_s_* ∑*_t_* ∑*_i_*(*C_ist_* + *U_st_*), where the summations apply only to cells (*s*) surveyed in Russian transects (i.e. the mean proportion of seals identified to species).

We assumed that counts (i.e., *C_ist_*, *U_st_*) followed a compound Poisson-Gamma (CPG) distribution with mean *μ_ist_*. The CPG distribution is a version of the Tweedie distribution where the power parameter (*ρ*) is limited to the range 1 < *ρ* < 2 (Jørgensen 1987). This three-parameter distribution is flexible, accommodating both zero inflation and overdispersion, and multiple authors have shown it to be appropriate for marine mammal count data (Miller et al. 2013; Sigourney et al. 2020). Separate power and dispersion parameters were estimated for each observation type (i.e. ringed, bearded, or unknown species counts) and survey platform (i.e. U.S. vs. Russian aircraft).

### 2.7 Model fitting and computing

We used maximum marginal likelihood to fit our model, with a joint pseudo-likelihood that included components for (1) U.S. and Russian count data, (2) prior distributions for regression parameters (including spline parameters), and (3) a penalty that forced the proportion of ringed seals basking after MDSDA melt dates to be 1.0. We used a parametric bootstrap procedure to propagate detection uncertainty into abundance estimates. For more detail on likelihood components, penalties, and bootstrap procedures see Supplement section 1. We coded our joint pseudo-likelihood function using Template Model Builder (TMB; Kristensen et al. 2016) syntax, and used the ‘nlminb’ function in R (R Core Team 2017) to optimize the likelihood. Importantly, TMB allowed us to use the Laplace approximation to integrate over prior distributions of smooth effects (which may be interpreted as random effects). We used randomized quantile residuals (RQR; Dunn and Smyth 1996) to assess goodness-of-fit (Supplement section 1). We compiled a standalone R package, ‘CHESSseal,’ that includes all data, R scripts, and TMB code needed to reproduce our analyses. It is available on GitHub at https://github.com/pconn/CHESSseal and will be published and archived at a long-term, publicly available repository upon manuscript acceptance.

### 2.8 Comparison to historical seal density estimates

Bengtson et al. (2005) estimated eastern Chukchi Sea ringed seal abundance (partially corrected for availability) and bearded seal density (not corrected for availability) from aerial surveys in 1999 and 2000. To compare our estimates to theirs, we attempted to match nearshore and offshore strata used in their surveys (Figure 1). We also used the mean availability predictions for our flights in late May to correct strata-specific Bengtson et al. (2005) bearded seal density estimates, reconstructing a survey-area-wide abundance estimate as

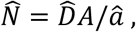

where 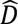 represents a density estimate, *A* is the area (km^2^) of sea ice habitat, and 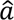 is the mean availability prediction. The surveys reported in Bengtson et al. (2005) occurred between May 21 and June 6. To compare our estimates to theirs, we summed predicted abundance for grid cells with centroids overlapping their survey areas. Because our procedure results in an estimate for each day of our survey, we took the mean value from May 21 through the end of our survey on May 31. We used the coefficient of variation from our previous bootstrap procedure to construct log-based confidence intervals (Burnham et al. 1987; Buckland et al. 2001) for our estimates. We used the delta method (Dorfman 1938) to calculate standard errors for abundance estimates associated with Bengtson et al. (2005) surveys.

Although there were surveys of seals in the Chukchi and Beaufort seas prior to 1999 (e.g., Frost et al. 1988; Frost et al. 2004), we do not compare our results to those surveys because those estimates were uncorrected for availability and perception probability.

### 2.9 Ethics approval

This research was conducted in conformance with all U.S., Russian Federation, and Canadian laws applicable to ringed and bearded seals. Alaska Fisheries Science Center (AFSC) conducted aerial surveys and deployed bio-loggers on bearded seals under National Marine Fisheries Service (NMFS) Research Permits 782-1765 and 15126, and Letters of Assurance of Compliance with Animal Welfare Act regulations, A/NW 2010-3 and A/NW 2016-1 from the AFSC/Northwest Fisheries Science Center Institutional Animal Care and Use Committee. Ringed and bearded seals were tagged with bio-loggers by Alaska Department of Fish and Game (ADF&G) and North Slope Borough Department of Wildlife Management (NSB-DWM) under NMFS Research Permits 358-1787-01, 15324, and 30466 issued to ADF&G, and under approved protocols by the ADF&G Animal Care and Use Committee 2006-16, 2014–03, 2015-25, 2016-23, 0027-2017-27, and 0027-2018-29. Ringed seal tagging with bio-loggers in Canada was conducted under Department of Fisheries and Oceans Scientific License number SLE-04/05-328 and SLE-05/06-322, and Animal Care and Use Protocol UFWI-ACC-2004-2005-001U.

## 3 Results

The U.S. survey aircraft completed 13,685 km of track line on survey effort between 7 April and 31 May 2016; the Russian Federation aircraft completed 13,961 km during 19–26 April and 12–18 May (Figure 1). The total area over which ringed and bearded seal abundances were estimated was 846,250 km^2^, 35% within the U.S. EEZ and 65% within the Russian Federation EEZ. The total numbers of individual seals photographed were 4,764 ringed seals, 1,094 bearded seals, and 180 unknown species. A total of 5,720 seals were observed in US waters, while 318 were observed in Russian waters. This difference in encounter rate was mostly due to differences in seal densities, although the Russian aircraft also had a lower overall detection probability for ringed seals due to higher behavioral response to the aircraft and fewer seals predicted to be basking during the Russian survey flights. For instance, the mean predicted detection probability for ringed seals was 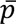 = 0.24 and 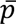 = 0.12 for U.S. and Russian surveys, respectively, but for bearded seals it was 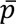 = 0.23 for each country.

We estimated that abundances of ringed and bearded seals in the Chukchi Sea were 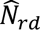 = 592,577 (95% CI: 478,448–733,929) and 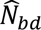 = 147,421 (95% CI: 114,155–190,380), respectively. Over the course of the study, the highest densities of ringed seals were observed in landfast ice in and around Kotzebue Sound, Alaska (Figure 2; see place names in Figure 1). Bearded seal densities were highest in offshore pack ice just north of the Bering Strait (Figure 2). Although the distribution of seals varied somewhat as ice conditions changed, time-averaged estimates of abundance suggested that 80% of ringed seals (point estimate 474,908) and 64% of bearded seals (point estimate 93,766) were in U.S. waters. The corresponding numbers for Russian waters, 117,669 ringed seals and 53,655 bearded seals, differ from those obtained by Chernook et al. (2019) because that initial analysis did not account for availability, among several other analytical differences. P-values from RQR goodness-of-fit tests were all greater than 0.10 for ringed seals, bearded seals, and unknown seal species in both U.S. and Russian surveys, suggesting that our models fit the count data reasonably well (Supplement section 1).

**Figure 2.**
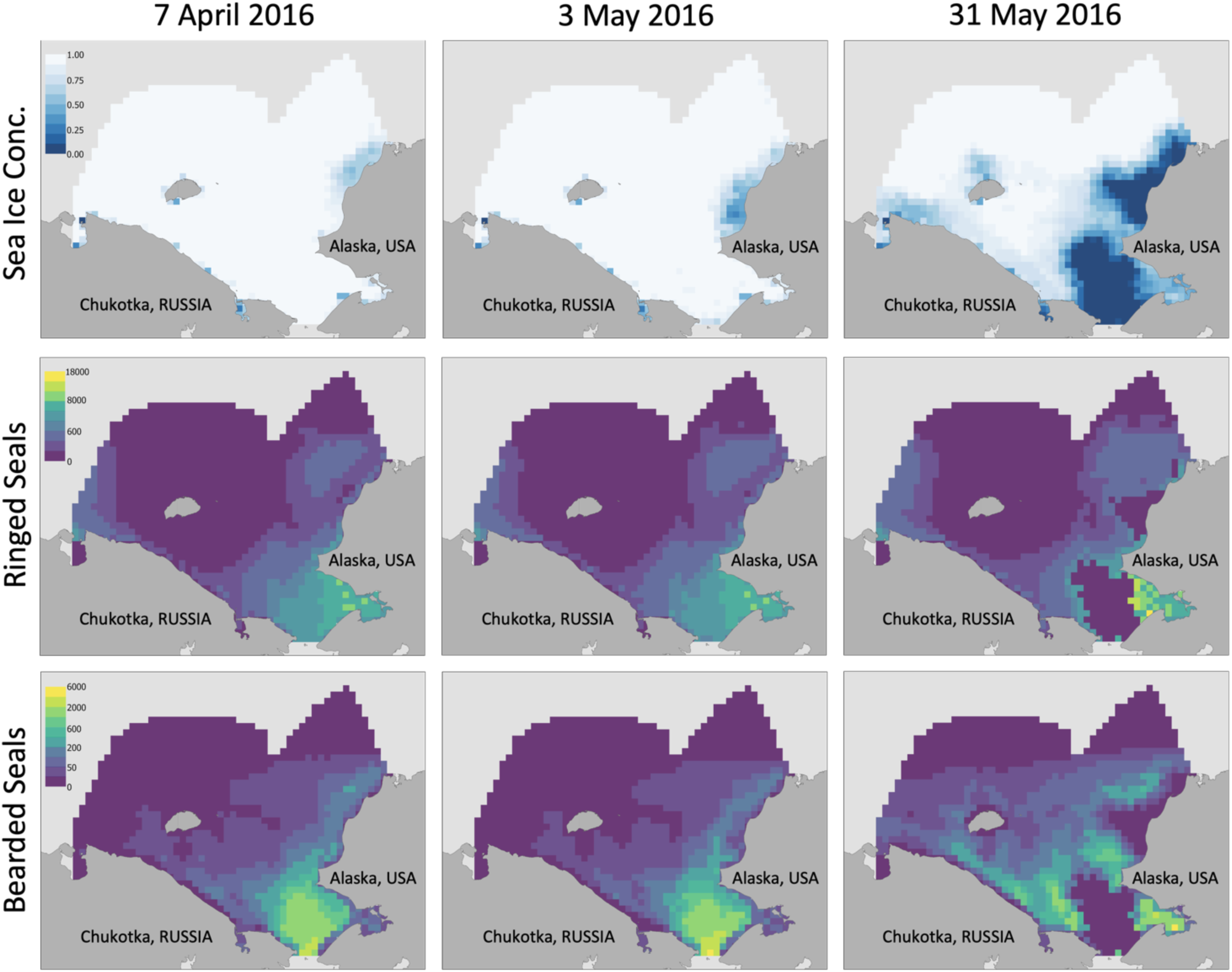
Remotely sensed sea ice concentration and estimates of ringed and bearded seal abundance (individuals per 25 × 25-km grid cell) in the Chukchi Sea on three dates at the beginning, middle, and end of the April–May, 2016 aerial surveys. Each column represents a different survey date, while rows correspond to sea ice concentration, ringed seal abundance, and bearded seal abundance. Note that the color scaling for seal abundance is nonlinear, depicting densities of zero in the darkest (indigo) color tone and very low densities in the second tone.

Ringed seal densities were dependent on northing and easting predictors—latitude and longitude effects, respectively—with densities (Figure 3) predicted to be the highest in the southeast quadrant of our study area, near Kotzebue Sound, and in the western edge of our study area, in the East Siberian Sea near Chaunskaya Bay, Chukotka (Figure 2). Densities increased slightly with sea ice concentrations above ≈0.6, but there was also a negative relationship (and high uncertainty) with lower ice concentrations. Ringed seal densities were high in cells with small proportions of landfast ice (<0.25) and in cells with intermediate to high proportions of landfast ice (>0.5), but lower in between these values (Figure 3). It should be noted that grid cells along much of the Alaska coastline (e.g. in locations north of Kotzebue Sound; Figure 1) had small proportions of landfast ice, simply because the size of the grid cells (≈625 km^2^) was large relative to the typically narrow band of landfast ice in each cell (Mahoney et al. 2014). Also, predicted ringed seal densities were high in Kotzebue Sound, where many cells had high or complete landfast ice cover. Densities increased strongly with snow depth values up to 20 cm, then reached a plateau between 20 and 30 cm (Figure 3). The modelled probability that a ringed seal was visible on the snow or ice surface (i.e. not in a lair), given that it was out of the water, increased from 0 at the beginning of the survey to 0.25 at about 15 days prior to the onset of snow melt indicated by the MDSDA index, (Figure 4); this probability then increased rapidly to 1.0 by about 5 days after snow melt onset.

**Figure 3.**
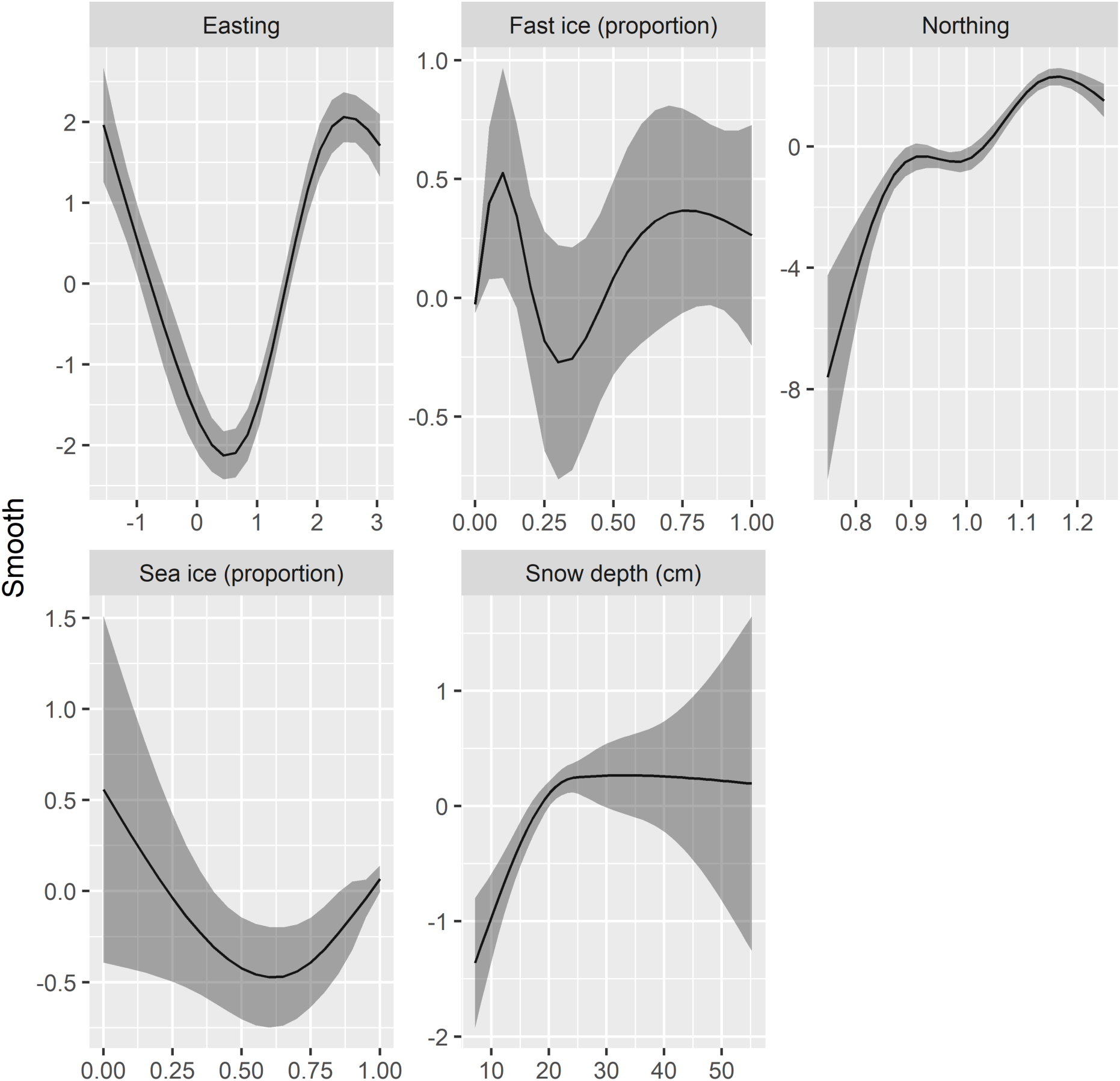
Estimated smooth functions (dark lines) for the relationship of ringed seal abundance to environmental and physiographic variables, on the multinomial logit scale, together with 95% confidence intervals (shaded areas). Note that the northing variable has larger values for locations farther south. Northing and easting variables were standardized to their means (so that a value of 1.0 roughly corresponds to the centroid of the study area). Landfast ice and sea ice values are proportions of area covered.

**Figure 4.**
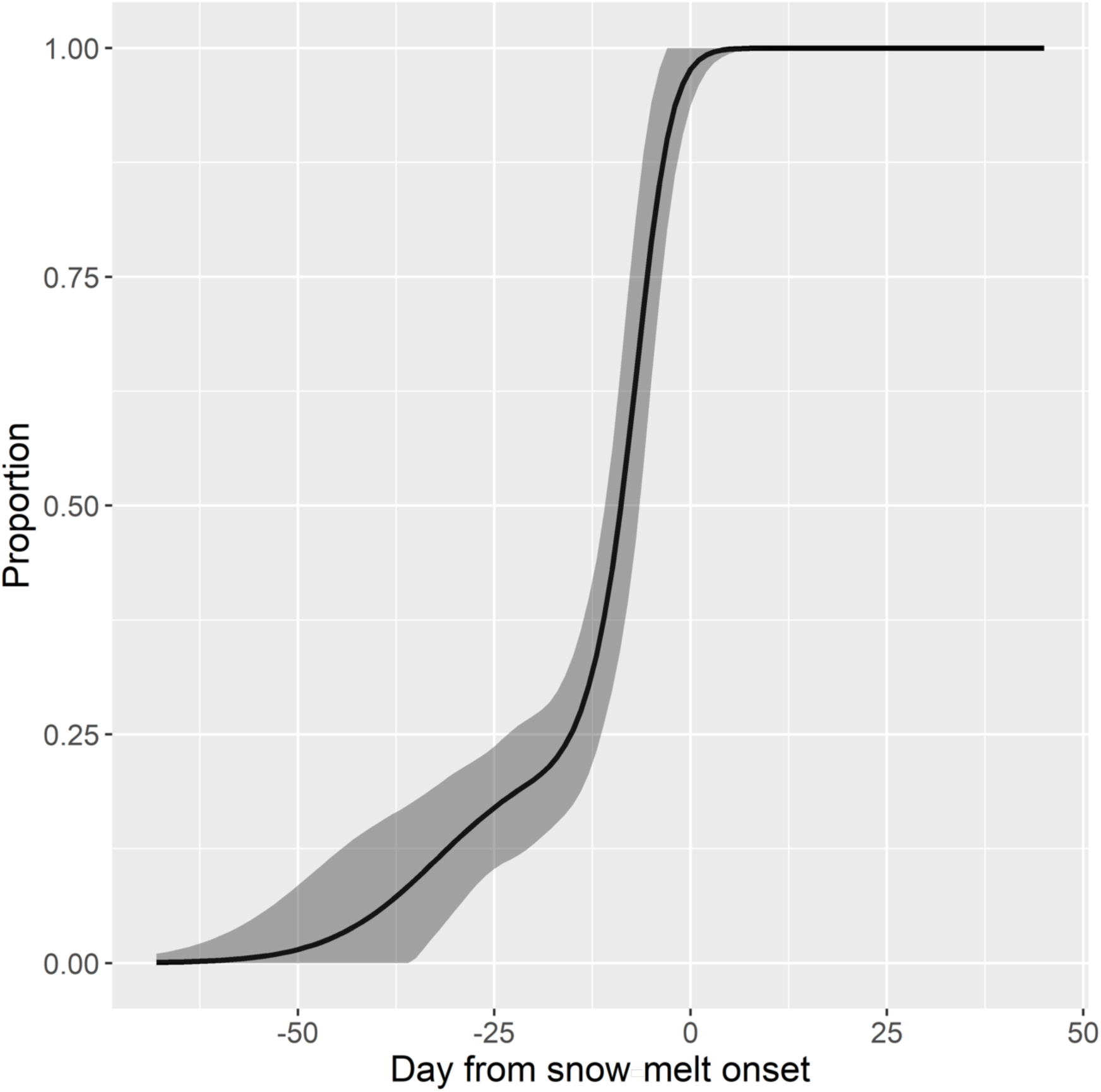
Proportion of ringed seals estimated to be basking on ice (i.e. not hidden in a subnivean lair) given that they were hauled out while aerial surveys were being conducted, together with 95% confidence intervals (shaded area). The model was penalized if the curve did not reach 1.0 by the date on which the MDSDA algorithm indicated the snow was saturated (‘snow melt onset’, 0 on x-axis).

The relationships between bearded seal abundance and predictive covariates (Figure 5) suggested that bearded seal densities were highest in grid cells without landfast ice, in cells with sea ice concentration near 0.8, and in cells with low snow depths. However, the most important predictor (as judged from the magnitude of y-axis value changes) was northing. Along with easting, these two covariates do not imply specific drivers of distributions. They may, however, highlight areas of low or high abundance that are not well predicted by the other, presumably more ecologically relevant covariates such as sea ice concentration.

**Figure 5.**
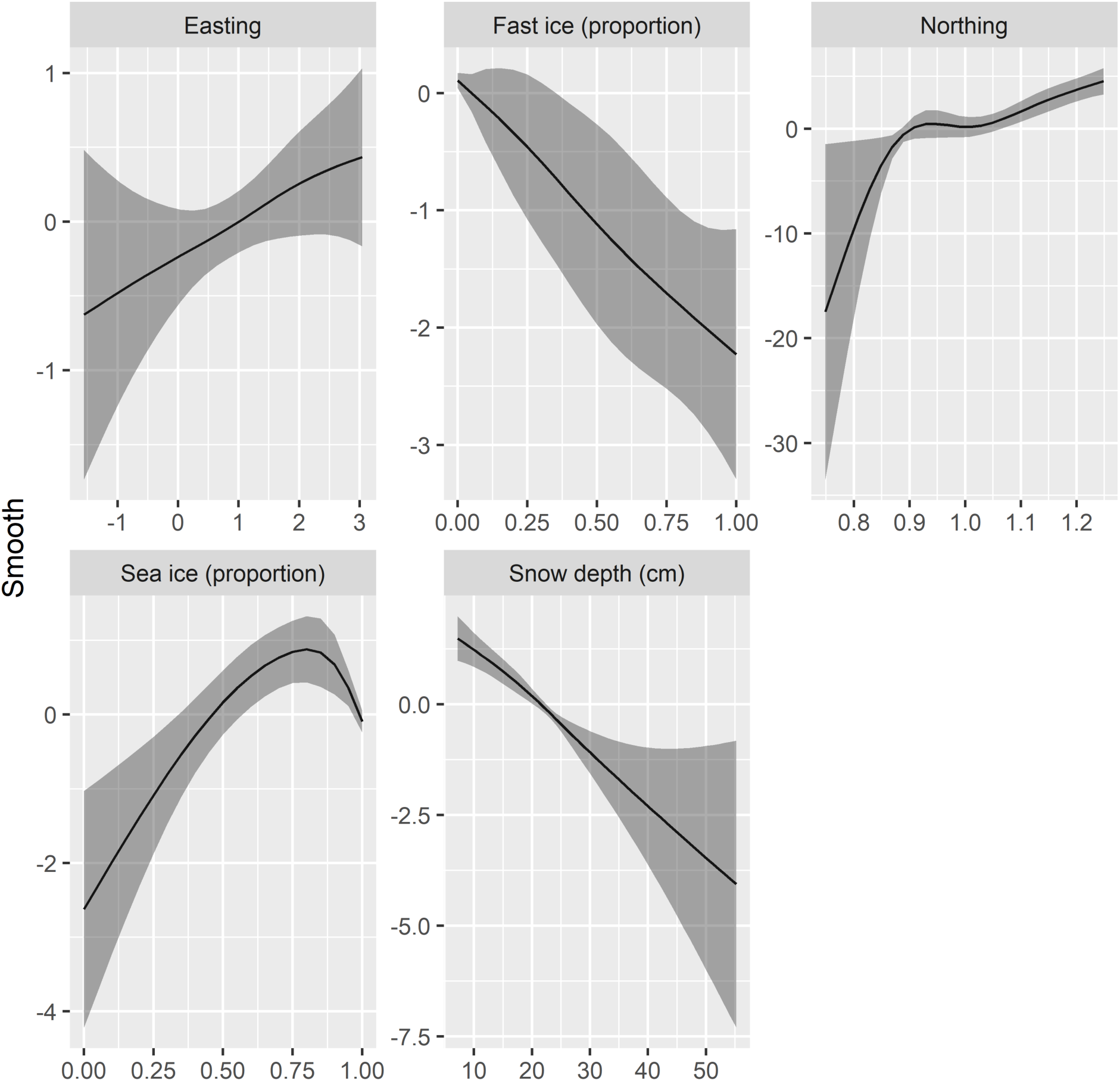
Estimated smooth functions (dark lines) for the relationship of bearded seal abundance to environmental and physiographic variables, on the multinomial logit scale, together with 95% confidence intervals (shaded areas). Note that the northing variable has larger values for locations farther south. Northing and easting variables were standardized to their means (so that a value of 1.0 roughly corresponds to the centroid of the study area). Landfast ice and sea ice values are proportions of area covered.

### 3.1 Comparison with historical estimates

Ringed seal abundance estimates reported for 1999 and 2000 in the eastern Chukchi Sea (blue shaded area in Figure 1) were 252,488 (SE 47,204) and 208,857 (SE 25,502), respectively (Bengtson et al. 2005). We re-constructed 95% log-based confidence intervals for those estimates as (175,576–363,091) and (164,550–265,095). For comparison, our ringed seal abundance estimate from the 2016 survey, in just the Bengtson et al. (2005) survey region (blue shaded area in Figure 1) at the end of May, was 394,411 (95% log-based CI: 318,449–448,493), about 1.7 times the average of the earlier estimates.

Bearded seal density estimates (Bengtson et al. 2005), adjusted with our availability estimates for late May (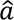 = 0.36, SE 0.03) led to abundance estimates of 25,076 (95% CI: 7,480–84,073) and 50,992 (95% CI: 30,207–86,079) for 1999 and 2000, respectively. Restricting abundance estimates from our 2016 surveys to grid cells with centroids in the Bengtson et al. (2005) study area (blue shaded area in Figure 1) led to a point estimate for late May of 72,182 (95% CI: 55,894–93,216), about 1.9 times the average of the earlier estimates.

## 4 Discussion

Our effort was the first to survey ringed and bearded seals over the entire Chukchi Sea, which was only possible through international cooperation. Conducting a sensor-based survey (as opposed to a survey with human observers) allowed us to fly faster and at higher altitudes, increasing the aircraft range and decreasing seal disturbance. We also used auxiliary data from bio-loggers to estimate availability, and employed protocols that allowed us to directly estimate the perception probability of our thermal sensing method and the proportion of seals that fled into the water before they could be detected. These are all improvements over previous surveys in the Chukchi Sea, which were limited in spatial scope and were able to only partially address perception and availability bias (e.g., Frost et al. 1988; Frost et al. 2004; Bengtson et al. 2005). In addition to providing a better abundance estimate, the results improve our understanding of ringed and bearded seal habitat composition, ecology, and prospects for persistence in rapidly warming Arctic marine ecosystems.

### 4.1 Factors influencing ringed and bearded seal distribution

Ringed seals were primarily found in the southern Chukchi Sea, concentrated in landfast ice in and around Kotzebue Sound; there were more moderate ringed seal densities in landfast ice elsewhere, along the northwest coast of Alaska and northeast coast of Russia near Chaunskaya Bay, Chukotka. Bearded seals also were primarily found in the southern Chukchi Sea, occupying broken pack ice with leads, similar to where they had been observed in past surveys. The broad geographic distribution patterns for both species in the U.S. portion of the Chukchi Sea match those found during surveys in 1999 and 2000 (Bengtson et al. 2005).

#### 4.1.1 Ringed seals

##### Influence of snow depth on ringed seal density

We found that ringed seal densities in spring were related to remotely sensed snow depth values, consistent with *in situ* studies of snow depth and the distribution of lairs. Resting lairs—also called simple, or haul-out lairs—are single-chambered structures used mostly by adults. Pups are typically found in multi-chambered structures—called pupping, pup, or birth lairs—that may include small tunnels excavated by the pups during their rearing period (e.g., Kelly et al. 1986). An analysis of ringed seal structures surveyed along the Alaskan Chukchi Sea coast during the 1980s found that the minimum snow depth at intact pupping lairs was 37 cm at Ledyard Bay and 58 cm in Kotzebue Sound, and the average depth at pupping lairs over the two sites was 75 cm (Hauser et al. 2021). Kelly et al. (2010b) reviewed other studies from Alaska, Canada, Russia, and Svalbard in which snow depths were measured at different types of ringed seal structures. Snow drifts to depths of 45 cm or more were found at resting lairs, and at least 50 cm—or more commonly, over 65 cm— were found at pupping lairs. Kelly et al. (2010b) concluded that average snow depths on flat ice of at least 20 to 30 cm are required to form drifts of adequate depths for excavation of lairs along ice deformations. In our results, ringed seal densities increased strongly with snow depth values up to 20 cm and plateaued between 20 and 30 cm, in close concurrence with the *in situ* findings that less than 20 cm average snow depth is suboptimal for viable lairs. The broad confidence intervals for ringed seal densities at average snow depths greater than 25 cm may reflect that the seals use areas of greater snow depths but are not necessarily found at high densities in all cells with deep snow; other habitat dimensions are also likely to be important. However, the narrow intervals and strong increase in seal densities as average snow depths increased up to 25 cm seem supportive of ringed seals’ selection of breeding habitat with sufficient snow for lair construction and integrity.

Remotely sensed snow depth products that provide information over the entire range of ringed seal habitat have only recently been tested for consistency over broad regions with the average snow depth requirements for lairs as derived from local, *in situ* studies. Lindsay et al. (2021) related the NASA Cryosphere database of 25-km gridded snow depths on sea ice (Markus and Cavalieri 1998; Comiso et al. 2003) to the raw counts of ringed seals detected in our surveys. There were modest but imprecise increases in counts of seals as with snow depths increased to about 25 cm for pups, and 35 cm for all ages combined. Lindsay et al. (2023) used an uncrewed aircraft to conduct photographic surveys of ringed seals on landfast ice near Kotzebue, Alaska in the spring of 2019. By relating seal counts to Landsat 8 210-m gridded pixel brightness values (an index of surface roughness) from May 6, 2019, and relating the brightness values to *in situ* April snow depths, they found indirectly that the highest seal counts occurred in areas where average April snow depth was 25 cm. In the current study, we related our ringed seal density estimates, rather than raw counts alone, to the NASA snow depth index, an estimate of the average depth over each 25 × 25-km grid cell. Our finding that ringed seal densities increased strongly with remotely sensed average snow depth up to 25 cm supports the conclusion from *in situ* measurements for this important habitat feature across the Chukchi Sea, an extensive region of the Pacific Arctic.

The capability to remotely monitor a snow-depth index of lair habitat quality over broad regions of the Arctic ringed seal’s range is likely to be helpful for assessing the subspecies’ status in the warming climate. A more complete understanding of lair habitat though, would include consideration of the timing and extent of sea ice formation, and patterns in ice surface structure, wind, air temperature, and precipitation. These factors interact to influence the blowing snow flux and the drift formation in the lee of ice surface structures (Kovacs et al. 2024); they are likely to vary among broad regions such as the Bering, Chukchi, and Beaufort Seas. We suspect that additional examples of aerial surveys for ringed seals, coupled with remotely sensed snow depths, may reveal regional variation in the relationship between ringed seal density and average snow depth alone.

##### Possible influence of covariates not in our model

Our results include the geographical covariates easting and northing that may identify low or high abundance areas that are not well predicted by other, presumably more ecologically relevant covariates in the model. They may also reflect influential covariates that are missing. We had no data to represent the distributions of ringed or bearded seal prey, which could influence seals’ distribution and density. We note, though, that the high densities of ringed seals we observed in Kotzebue Sound are a regular winter and spring occurrence reported by Indigenous hunters (Lindsay et al. 2023) that may reflect dependable winter and spring prey availability. Saffron cod (*Eleginus gracilis*) is highly prevalent in Alaska ringed seal diets (Quakenbush et al. 2020). Although saffron cod spawning grounds are not well known in Alaska, Kotzebue Sound and the adjacent southeast Chukchi Sea include large areas of shallow, nearshore waters that are typically ice covered from December to March—consistent with saffron cod spawning habitat (Wolotira 1985; Vestfals et al. 2019). Using survey catches of saffron cod larvae in the Chukchi Sea, coupled with biophysical transport models, Deary et al. (2021) suggested that Kotzebue Sound is an important saffron cod spawning ground. With the peak of spawning occurring in winter, gravid saffron cod would be ideal prey for ringed seals, especially pregnant females, which give birth to pups in March and April. The combination of dependable seasonal ice cover—much of which is landfast—and a reliable prey base could explain the high densities of ringed seals observed in Kotzebue Sound and southeast Chukchi Sea surveys over several decades (Frost et al. 1988; Bengtson et al. 2005, this study), and the reliance of Indigenous peoples of the region for winter sustenance over millennia (Lucier and VanStone 1991).

#### 4.1.2 Bearded seals

In past aerial surveys during March–May, primarily in the Bering Sea, bearded seal densities were highest in sea ice concentrations greater than 25%. There were density peaks at concentrations of 70–90% (Simpkins et al. 2003) and 67% (Conn et al. 2014), though Ver Hoef et al. (2014) found only a weak increase in the probability of occurrence in concentrations above 25% and no evidence for an effect of ice concentration on bearded seal density. A sample of juvenile bearded seals tagged in 2004–2009 and tracked during spring in the Bering Sea seemed to prefer ice concentrations similar to those identified from aerial surveys (∼80%; Cameron et al. 2018) or lower (∼50–60%; Breed et al. 2018). For juvenile bearded seals tagged in a more recent and warmer period of 2014–2018, there was only weak selection by juveniles in spring, for ice concentration <50%, perhaps reflecting a shift in the distribution of ice concentrations or an indication that other habitat features such as foraging quality are more important (Olnes et al. 2021). Our finding of a strong increase in bearded seal densities with sea ice concentration up to a peak at 80%, and a slight decrease thereafter, agrees more closely with previous aerial survey results (based on all age classes) than with results from tracking juveniles. Together with the strong decline we observed in bearded seal density with increasing proportions of landfast ice in a survey grid cell, our results are also consistent with the many qualitative natural history descriptions of bearded seals’ preference for broken, drifting ice, as well as their typical (but not complete) avoidance of the heaviest concentrations of pack ice and extensive landfast ice (Fedoseev 1965; Burns and Harbo 1977; Burns and Frost 1979; Burns 1981; Smith 1981; Fedoseev 1984; Nelson et al. 1984; Kingsley et al. 1985; Cameron et al. 2010). While we also observed a decline in bearded seal density with increasing snow depth, we are not aware of prior studies documenting a negative association with snow depth. Lacking an ecological explanation based on bearded seal natural history, this result may serve as a reminder that models with multiple covariates sometimes yield illusory associations. For example, if there is a latitudinal (northing) gradient in snow depth, these two variables would not be independent and thus difficult to interpret. Deeper snow depth over areas of landfast ice could also contribute to the apparent association between bearded seal densities and areas of shallower snow depth.

### 4.2 Assumptions and limitations

We modeled ringed seal availability as a function of a snow melt index (i.e., Belchansky et al. 2004) that was previously found to be related to an increase in the proportion of seals basking (Kelly et al. 2006). Our model could accommodate a gradual transition away from lair use—including intermittent lair use by individuals—but it could not account for a portion of ringed seals that may continue to spend time in lairs after the snow melt onset. We assumed that all individuals transitioned to basking by the date of snow melt onset or shortly thereafter, and we had no means to assess any negative bias associated with this assumption. Kelly et al. (2006) found strong correlations between the end of lair use by tagged ringed seals in the Beaufort Sea and the dates of melt onset from the MDSDA and another snow melt index. If those correlations are applicable to our study, we would expect the bias from our assumption to be low, but the sample size (*n*=5 years) was small and the correlations weren’t adjusted for testing multiple snow melt indices. We also had no means to assess imprecision associated with this assumption, and we caution against interpreting our confidence intervals too rigidly; they are likely too narrow for ringed seals despite having included several more sources of uncertainty in our model than previous examples of aerial surveys for seals. Future research should be devoted to refining ringed seal availability estimates through analysis of existing data (e.g. Lindsay et al. 2021) and field work (e.g. applying additional bio-loggers, investigating relationships between internal lair conditions and snow melt onset, and examining variations in emergence timing as a function of environmental covariates).

The timing of our surveys of the Chukchi Sea in 2016 was, in part, a result of our desire to combine Chukchi Sea estimates with Bering Sea estimates from a previous survey (e.g. Conn et al. 2014) and estimates from surveys planned for the Beaufort Sea, in order to develop total estimates for the Beringia DPS of bearded seals and for ringed and bearded seals in U.S. waters (i.e. the Alaska stocks for assessment under the Marine Mammal Protection Act; Young et al. 2023). In particular, we wanted to avoid surveying in June when large numbers of seals migrate north from the Bering Sea to the Chukchi Sea (Melnikov 2017; Melnikov 2022). However, this approach may not have been the best timing for ringed seals, as VHF radio-tagging data and monitoring of ringed seal lairs in the Beaufort Sea (Kelly et al. 2006; Kelly et al. 2010a; Kelly et al. 2010b) suggest that a large proportion of seals were in lairs and thus were unavailable for counting when our surveys began in early April. Smith (1981) suggested that optimal timing of bearded seal surveys might be late April–May when mothers are accompanied by pups, at least a month earlier than the best time for ringed seal surveys. If ringed seals are of higher priority in future surveys, we suggest that more intensive surveys in the latter half of May and early June may be more useful, as it would remove a large source of uncertainty about availability in April and early May. Also, aircraft capable of surveying at sufficient speed could allow coverage of both the Bering and Chukchi seas in a single survey period, reducing the potential for bias induced by seals migrating north during the survey.

We avoided surveying small portions of the study area that occurred within 48 km (30 miles) of the Alaskan villages of Utqiaġvik, Wainwright, Point Lay, and Point Hope, to reduce the potential for disturbance of traditional bowhead whale (*Balaena mysticetus*) harvest activities. Coastal Indigenous village sites in the Arctic are recognized as places of dependable, high densities of marine resources. Therefore, not surveying near these villages could potentially have biased our estimates low. Our analysis estimated that 4% of the totals for both ringed and bearded seals occurred within these avoided areas, which composed 2.1% of the total study area. Therefore, because our estimated seal densities in these avoided areas were roughly twice the overall average for each species, and because these areas were very small in comparison to the total study area, we consider that excluding them was unlikely to have caused significant bias in our overall abundance estimates. Nevertheless, it would be beneficial in future surveys to incorporate survey effort in those areas if it can be done in coordination with the communities to avoid interfering with traditional subsistence activities.

### 4.3 Implications for future status of ringed and bearded seals

#### Relationships between snow depth and ringed seal vital rates

Several studies have found associations between regional snow depth and ringed seal survival, reproduction, or indices of condition (e.g. see review by Kelly et al. 2010b). Hammill and Smith (1991) found that a decrease from 23 cm to 10 cm in average snow depth in Barrow Strait, Canada in the mid-1980s coincided with an increase in the predation rate by polar bears from 0.1 to 0.4 seals/km^2^. An index of recruitment from seals harvested in western Hudson Bay during 1991–2001 was sharply lower for cohorts born in years with spring average snow depths less than about 32 cm (Ferguson et al. 2005). Chambellant et al. (2012) found that an index of pup survival was optimized between about 25 and 52 cm of snow depth (peak at 38 cm) measured on the ground near their study area in western Hudson Bay during 1991–2006. Iacozza and Ferguson (2014) found that variability in remotely sensed spring snow depth in western Hudson Bay from 2003–2010 was negatively related to the proportion of young-of-the-year in the autumn harvest at Arviat, Canada; the number of ‘dry snow cover days’, which is an index of the duration of winter snow cover, was positively related to pup survival, and together, the two indices accounted for 82% of the variation in the young-of-the-year harvest numbers. Hamilton et al. (2018) found that ringed seals hauled out less frequently and—during winter—in shorter bouts and for less time overall, after the cessation of annual formation of landfast ice that occurred in 2006 around Svalbard. Although the changes were not directly linked to vital rates, the implication was that without landfast ice and its associated snow cover, ringed seals hauled out less in lairs, with likely impacts on their energy budgets and predation risk. But also in Svalbard, where large areas have not had regular seasonal sea ice since 2006, an interdecadal synthesis of several vital rate indices found no clear pattern consistent with climate-related hypotheses (Andersen et al. 2021). Potential explanations are ‘life history plasticity’ or that small-scale variations in breeding habitat remain suitable in the limited area where the hunted samples were collected (Andersen et al. 2021).

Around Alaska and the western Canadian Arctic, associations among ringed seal vital rates or condition and snow depth on sea ice have not been assessed, though some studies have made comparisons to ice-covered area, the length of the ice-covered period, or other statistics that may be related to spring snow depth. Crawford et al. (2015) concluded that young ringed seals in the Chukchi and northern Bering seas grew faster during 2003–2012, which was characterized as a period of reduced sea ice, compared to an earlier period with more ice, 1974–1984. Ringed seals harvested at three northern Bering Sea and one southern Chukchi Sea villages included higher proportions of pups during the recent period, as well. A continuation of the study, comparing the period 2000–2015 with 2016–2020 (Quakenbush et al. 2020), found that growth of ringed seals born in the most recent period did not vary from average, except those born in 2017. Blubber thickness (i.e. body condition) was below average in 2017–2018 but returned to average in 2019. The years of 2017 and 2018 were extremely warm in the Pacific Arctic, with low sea ice coverage and impacts documented throughout the ecosystem (Huntington et al. 2020).

In the western Canadian Arctic, ringed seals were monitored in the Amundsen Gulf and western Prince Albert Sound during 36 years between 1971 and 2019 (Harwood et al. 2012; Harwood et al. 2020). These areas were characterized as prime habitat, with sea ice that typically persists well beyond the ringed seal pup rearing period. Despite long-term trends toward later autumn freeze-up and earlier spring sea ice clearance that coincided with a declining trend in seal body condition, the poorest body condition and lowest indices of reproduction occurred in the years with the latest ice break-up, suggesting more complex processes than a simple directional influence of Arctic warming on the seals’ vital rates.

In general, the studies summarized above have examined suites of indices for vital rates but have not had the means to discern overall demographic impacts, such as changes in abundance or loss of significant portions of breeding range. Although monitoring indices of vital rates or other demographic parameters provides insight about mechanisms of response to environmental variability and climatic trends, sampling considerations are often complex, as are interpretations of the results; the need remains for complementary, direct means of monitoring abundance and distribution of breeding populations, such as count-based surveys.

#### Trends in snow depth on sea ice

Snow depth on Arctic sea ice has decreased, prompting the question of whether ringed seal populations have already been impacted by diminished or degraded habitat. Warming in the upper layer of Arctic waters has caused later onset of sea ice formation in autumn, which in turn reduces snow accumulation on the ice because more snow falls into open water (Webster et al. 2018). The result has been a clear multidecadal decline in average snow depth on sea ice. For example, Webster et al. (2014) found that snow on Chukchi and Beaufort sea ice in 2009–2013 was about 56% thinner than during the latter half of the 20^th^ century. March–May snow depth on sea ice at three long-term monitoring sites in the Canadian Arctic declined by 0.1, 1.7, and 0.6 cm/decade from 1960–2020 (but increased by 0.1 cm/decade at a fourth site; Lam et al. 2022). Lee et al. (2021) found an Arctic-wide average reduction in snow depth from the period 1954–1991 to the period 2003–2020, but with a statistically significant positive trend of ∼0.6 cm/year on multi-year ice, and negative trends (∼−0.4 cm/year) over mixed and first-year ice, the habitat for most Arctic ringed seals. Stroeve et al. (2020), using a detailed physical model of snow depth driven by weather reanalysis products, estimated a loss of 1.9–3.0 cm/decade from 1980–2016 in the Chukchi Sea. A version of the same model was used to estimate declining trends over the past 2–3 decades in snow depth, blowing snow flux, and ringed seal lair habitat around Svalbard (Kovacs et al. 2024). Dou et al. (2021) showed that rain-on-snow events in the ERA5 reanalysis have shifted as much as 4–6 days earlier per decade in some regions; there have been more rain-on-snow events in spring; and the increased rain-on-snow events have caused a reduction in spring snow depth of 0.5 cm/decade over the Arctic Ocean since 1980 and up to 2 cm/decade in the Kara-Barents seas and Canadian Arctic Archipelago. The observed open-water period in the Chukchi Sea has increased by about 2 days/year over the period 1979–2013 (Crawford et al. 2021). The length of the open water period is particularly relevant to habitat quality for Arctic ringed seals because snow depth and quality is influenced by the ice freeze-up and melt onset dates.

#### Reference levels for climate change impacts on ringed and bearded seals in the Chukchi Sea

Given the observed and ongoing declines in Arctic sea ice and snow depth, do our estimates in 2016 provide a reference for assessing climate-driven population trends of ringed and bearded seals in the Chukchi Sea? The large numbers of both species that we estimated and the apparent increases since a previous, smaller scale survey are encouraging, especially given the current designations of both species as ‘Threatened’ under the U.S. Endangered Species Act (National Marine Fisheries Service 2012a; National Marine Fisheries Service 2012b). However, increases in Chukchi Sea ringed and bearded seal estimates could be due to factors other than population growth, such as shifting of breeding locales north from the Bering Sea, shifts to earlier northward spring migration from the Bering Sea, or differences in survey methods. For instance, although ringed seal densities reported by Bengtson et al. (2005) were adjusted for the haul-out component of availability, there was no correction for ringed seals that may have been hidden in snow lairs. The double-observer method they used to correct for incomplete perception is now recognized to be biased if assumptions of independence are not met, which is often the case in aerial surveys (e.g., Conn et al. 2012; Burt et al. 2014). Further, their estimates did not take into account disturbance of seals into the water by the aircraft, which flew at an altitude of 91 m, compared to altitudes of 250–300 m during our survey. All of these factors would negatively bias abundance estimators from Bengtson et al. (2005), and could plausibly account for most or all of the increases in abundance estimates (1.7 and 1.9 times greater for ringed and bearded seals, respectively). However, some contribution from actual changes in abundance or shifts distribution cannot be ruled out. We took steps to accommodate the known sources of detection bias in previous surveys, making future comparisons more straightforward.

The studies of vital rates that we summarized earlier indicate that Arctic ringed seals in some parts of their range—Hudson Bay and around Svalbard—have responded to years of poor snow cover on sea ice in ways that would be expected to impact vital rates and reduce population size under long-term Arctic warming. Both of those regions are peripheral, rather than core ringed seal habitat, due to low latitude (Hudson Bay) or oceanographic features that mediate Arctic conditions (Svalbard). In contrast, where ringed seals have been studied in the core areas of their range, they have yet to exhibit consistent negative impacts from climatic trends. In the Alaska Chukchi Sea and in the western Canadian Arctic, some of the most notable reductions in indices of vital rates and condition have actually come during colder periods or years with higher ice coverage and later ice breakup (Crawford et al. 2015; Harwood et al. 2020). We suggest that our spring 2016 estimate of ringed seal abundance in the Chukchi Sea represents a baseline reference level for assessing future climate-related changes in abundance. Similarly, bearded seals in Alaska have not exhibited declines in body condition, growth, or reproduction compared to earlier periods (Crawford et al. 2015; Quakenbush 2020), and they are not dependent upon snow cover for shelter of their young. We are not aware of other scientific evidence for climate-driven impacts on bearded seals of the Chukchi Sea, nor of impacts documented by Indigenous hunters and observers in the region. Therefore, we suggest that our bearded seal abundance estimate is also a reasonable reference point for assessment of future climate-related impacts to that species.

How ringed and bearded seals will respond to future climate change remains uncertain. Will seals migrate northward earlier as spring sea ice diminishes in the Bering Sea? Will there be enough prey of sufficient quality to accommodate them? Will snow depths on sea ice, which are predicted to continue declining over the next century (Hezel et al. 2012; Webster et al. 2021), be sufficient for ringed seal lair construction? Although current populations in the Chukchi Sea appear healthy, there is cause for concern about how they will cope with a rapidly warming Arctic. The upper and lower bounds of our 95% confidence intervals for ringed and bearded seal abundance were about 25% above and 20% below the point estimates, respectively. These levels of precision provide reasonable expectations for detecting substantial changes in Arctic seal populations over broad regions using methods like those we have demonstrated in the Chukchi Sea. For example, with a coefficient of variation (CV) of 13% that reflects the precision of our bearded seal estimate, a 25% decline in the population would have a 33% chance of being detected in a subsequent survey, whereas a 50% decline would be detected 97% of the time (95% two-sided z-test). Our ringed seal estimate, with a CV of 11%, was nominally more precise, but was based on additional assumptions that are not reflected in the precision estimate. We consider the two species to be similar for the purposes of detecting future trends. Increasing the number of subsequent surveys would increase the power to detect changes in a regression framework. Future instrument-based surveys in the Chukchi Sea and other areas (e.g. Bering and Beaufort seas) will help us measure ringed and bearded seals’ responses to the dramatic changes occurring in the Arctic.

## 5. Acknowledgements

We are grateful to Gavin Brady, Sarah Brown, Shawn Dahle, and Carter Johnson for species identification and detection rate estimation; John Bengtson for project facilitation as co-Convenor of the Working Group on Marine Mammals under Area V of the U.S.-Russia Agreement on Cooperation in the Field of Environmental Protection and Natural Resources; Capt. Robert Dodson, 1st Officer Sarah Morris and Aircraft Mechanic James DeLuca (“JD”) of Dynamic Aviation for safe operation of the King Air A90; John Cabrer of MoviTHERM for collaboration on Skeyes software and the hotspot detection method; the late Craig George for logistical support and coordination with whaling activities in Utqiaġvik; Paul Wade for helpful comments on the manuscript; Dave Anderson and Rich Lomire of the National Weather Service for assistance with housing; the U.S. Fish and Wildlife Service for providing cooled LWIR cameras; the Russian non-profit organization, Marine Mammal Council, provided logistical support for aerial surveys; N.A. Chernook and V.V. Asyutenko for processing Russian survey images; and D. M. Glazov for contribution to the field effort. The organization of this international collaboration would not have been possible without the facilitation by Vladimir N. Burkanov of North Pacific Wildlife Consultants.

The projects that deployed bio-loggers, overseen by ADF&G and NSB-DWM, were successful because of support and cooperation of the Ice Seal Committee, the NSB Fish and Game Management Committee, and coastal Native communities, especially the Native Village of Barrow, Native Village of Kotzebue, Native Village of Koyuk, and the Native Village of Saint Michael. The success of the studies was due to assistance from subsistence hunters—especially John Goodwin, Merlin Henry, Joe Skin, Isaac Leavitt, Bobby Sarren, and Billy Adams—and biologists, especially Mark Nelson, Anna Bryan, Ryan Adam, Rowenna Gryba, Aaron Morris, and Jason Herreman. Expertise and leadership in the field for tagging seals in Canadian waters was provided by Tom Smith; community crews from Tuktoyaktuk, Inuvik and Paulatuk, Northwest Territories; and Brendan Kelly and John Moran of the University of Alaska.

## 7. Author contributions

Conceptualization: Peter Boveng, Vladimir Chernook; Data curation: Stacie Koslovsky; Formal analysis: Paul Conn, Joshua London, Brett McClintock, Irina Trukhanova; Funding acquisition: Peter Boveng, Vladimir Chernook; Investigation: Vladimir Chernook, Erin Moreland, Paul Conn, Cynthia Christman, Benjamin Hou, Denis Litovka, Erin Richmond, Alexander Vasiliev, Amy Willoughby; Methodology: Peter Boveng, Vladimir Chernook, Paul Conn, Benjamin Hou, Jessica Lindsay, Erin Moreland, Alexander Vasiliev; Project administration: Michael Cameron, Erin Moreland; Resources: Justin Crawford, Lois Harwood, Nikita Platonov, Lori Quakenbush, Andrew Von Duyke; Software: Benjamin Hou, Stacie Koslovsky; Supervision: Peter Boveng, Vladimir Chernook; Validation: Paul Conn, Stacie Koslovsky, Erin Moreland; Visualization: Paul Conn, Joshua London, Erin Richmond; Writing - original draft: Peter Boveng, Paul Conn; Writing - review and editing: All authors.

## 9. Data Availability

All data and code will be made freely available as a GitHub repository upon acceptance for publication in a peer-reviewed journal.

