## Supplements_1_and_2 for "Abundance and distribution of ringed and bearded seals in the Chukchi Sea: a reference for future trends"

### 1. ADDITIONAL DETAILS FOR MODELING SEAL COUNTS

Here we provide additional information on (1) modeling habitat and landscape effects on seal densities (including p-splines), (2) construction of detection prior distributions, (3) construction of the joint marginal pseudo-likelihood used for estimation, (4) the parametric bootstrapping approach for propagating uncertainty in detection probability into abundance estimates, and (5) goodness-of-fit procedures and goodness-of-fit results.

#### 1.1. HABITAT COVARIATE MODELLING

As indicated in the main manuscript, our model computed time ($t$), location ($s$), and species-specific ($i$) abundance, $N_{ist}$, as

$$N_{ist} = N_i \frac{w_s \exp(\boldsymbol{x}_{ist}\boldsymbol{\beta})}{\sum_j w_j \exp(\boldsymbol{x}_{ijt}\boldsymbol{\beta})},$$

where $w_s$ is the proportion of grid cell $s$ that is composed of marine habitat (i.e. omitting land), $\boldsymbol{x}_{ist}$ is a row vector of explanatory variables for species $i$ in cell $s$ at time $t$, and $\boldsymbol{\beta}$ is a column vector of regression coefficients. One approach with such a model is to specify $\boldsymbol{x}_{ist}$ by using some known functional forms (e.g. linear and/or quadratic effects in the case of polynomial regression). However, we wanted additional flexibility and instead implemented a penalized spline formulation for habitat and landscape covariates, and a penalty on $\boldsymbol{\beta}$ parameters to prevent the model from being too flexible. Our approach is very much in the spirit of generalized additive models (Wood 2017) and largely followed the template for ‘psplines’ available at https://github.com/skaug/tmb-case-studies. In particular, we used the ‘gam’ function in the mgcv package (Wood 2017) in the R programming environment (R Development Core Team 2017) to construct cubic smoothing bases and penalty matrices for continuous covariates. Initial testing suggested some instability when the dimension of these bases was unconstrained, so we set this dimension as $k = 6$. The result is that $\boldsymbol{x}_{ist}$ includes six predictors for each continuous covariate.

Spline models are typically overparameterized in the sense that they will often result in relationships that are too “wiggly” without some sort of penalization of smooth terms. A spline penalization was thus included in the form of a set of prior distributions on $\boldsymbol{\beta}$ parameters. Without loss of generality, let us assume that $\boldsymbol{\beta}_1 = \{\beta_1, \beta_2, \cdots, \beta_6\}$ represents regression coefficients for the first smooth term. Then our prior distribution for $\boldsymbol{\beta}_1$ would be

$$[\boldsymbol{\beta}_1] \equiv \text{Multivariate normal}(0, (\lambda_1 \boldsymbol{S}_1)^{-1})$$

This is the formulation used by Wood (2017) and in his 'jagam' function included in the mgcv package (Wood 2011). We included the same gamma priors for the penalty coefficients, $\lambda_i$, used in the 'jagam' package:

$$[\lambda_i] \equiv \text{Gamma}(0.05, 0.005).$$

In addition to smooth terms, we included a fixed effect for the binary indicator 'I_no_ice.' Because seals cannot be detected in open water, we set the fixed effect equal to -20.0 on the linear predictor scale to force abundance to be close to zero in those cells.

#### 1.2. Detection priors

We decompose detection probability for species $i$ at time $t$ in cell $s$ as

$$p_{ist} = a_{ist} b_i d,$$

where $a_{ist}$ is availability, or the probability that a seal is basking on ice (i.e. on ice and visible to aircraft) during sampling, $b_i$ is a non-disturbance probability (i.e. the probability that a basking seal remains on ice and is not flushed into the water the aircraft flies overhead), and $d$ is the overall detection probability of the thermal sensors and a semi-automated detection process. To propagate uncertainty about these various components of detection into final abundance estimates, we used auxiliary data to produce a prior distribution for $p_{ist}$ on the logit scale.

To measure detection probability of thermal sensors, we used data from a double sampling study in which an unaided human analyst examined 28,052 color images from 2016 survey flights. In these photos they found 258 seals, and these "test" data were used to find the proportion of those seals that were detected using an algorithm applied to our thermal images. In total, we detected 247/258 seals in this manner, so that $\hat{d} = 0.96$, with Bernoulli variance $\widehat{Var}(\hat{d}) = 0.0002$. In absence of a similar study of the instruments used by our Russian Federation collaborators, we assumed the Russian sensors had the same detection probability.

To measure non-disturbance probability, $b_i$, we conducted forward-facing disturbance trials. In these trials, our pilot and copilot visually detected seals at long distances in front of the aircraft, before they had a chance to respond to aircraft (i.e. were not detected based on movement). We then recorded whether or not these seals flushed into the water before our aircraft passed overhead. For U.S. surveys, 6/29 ringed seals flushed into the water and 0/14 bearded seals flushed into the water. In Russian surveys, 57/189 ringed seals and 3/51 bearded seals flushed into the water. Letting $i = 1$ denote ringed seals and $i = 2$ denote bearded seals, we have $\hat{b}_1 = 0.79$, and $\widehat{\text{Var}}(\hat{b}_1) = 0.0057$ for U.S. surveys and $\hat{b}_1 = 0.70$, and $\widehat{\text{Var}}(\hat{b}_1) = 0.0011$ for Russian surveys. For bearded seals, $\hat{b}_2 = 1.0$, $\widehat{\text{Var}}(\hat{b}_2) = 0.00$ for U.S. surveys and have $\hat{b}_2 = 0.94$, $\widehat{\text{Var}}(\hat{b}_2) = 0.0011$ for Russian surveys.

We modeled availability differently for ringed and bearded seals. Prior to this study, ringed seal availability had not been previously analyzed outside of several studies with limited sample size (e.g. Bengtson et al. 2004, Von Duyke et al. 2020). Further, it was difficult to equate availability to haul-out proportions from TDRs affixed to ringed seals, particularly early in our surveys when ringed seals could be hauled out but still unavailable because they were hidden in subnivean lairs (Kelly et al. 2006). We thus conducted a standalone analysis to estimate haul-out proportions as well as changes in diel behavior that might indicate emergence from subnivean lairs (Section 2). As indicated in the main manuscript text, our general approach for ringed seals was to decompose availability ($a_{st}$) at time $t$ and location $s$ as $a_{st} = h_{st}\alpha_{st}$, where $h_{st}$ represents the probability a ringed seal is hauled out, and $\alpha_{st}$ represents the probability that a hauled-out ringed seal is visible to airborne sensors (i.e. is not in a subnivean lair). Section 2 describes how $h_{st}$ was estimated.

Our approach to modelling availability in ringed seals was to embed a flexible penalized spline model for $\alpha_{st}$, the proportion of hauled out seals who are also visible to aircraft, within the integrated model for survey counts. The model was penalized for differing from 1.0 survey dates after the snow melt onset date indicated by the MDSDA algorithm (see main text). Prior to the snow melt onset date, $\alpha_{st}$ was allowed to smoothly vary in a way that best represented temporal trends in survey counts. Our intuition was that the increase in survey counts throughout the course of the survey provides information on the relationship between availability and snow melt onset. However, such trends in counts do not provide information about the absolute level of availability, and thus the constraint of setting it to 1.0 after the melt date helped to determine the scale of the availability curve. In particular we modeled

$$p_{1st} = \frac{1}{1+\exp(-\nu_{1st})}\frac{1}{1+\exp(-\mathbf{y_{st}}\boldsymbol{\kappa})},$$

where $\mathbf{y_{st}}$ is a cubic spline basis vector for Julian day obtained from `mgcv` (with 7 degrees of freedom) evaluated at $\delta_{st}$, and $\boldsymbol{\kappa}$ is a vector of spline coefficients (note that $\alpha_{st} = [1 + \exp(-\mathbf{y_{st}}\boldsymbol{\kappa})]^{-1}$). Once again, $\nu_{1st}$ represented logit-scale detection effects using an informed prior:

$$[\boldsymbol{\nu}_1] \equiv \text{Multivariate normal}(\boldsymbol{\mu}_{1p}, \boldsymbol{\Sigma}_{1p}) . \qquad \text{Eq. S1}$$

In this case, our prior mean and variance computations represented a product of non-disturbance ($b$), detection of thermal sensors and the semi-automated processing algorithm ($d$), haul-out probability predictions ($\tilde{h}_{1st}$), with $\boldsymbol{\mu}_{1p} = \text{logit}(b_1\tilde{h}_{1st}d)$; we used the delta method to compute $\boldsymbol{\Sigma}_{1p}$.

For bearded seals, we used data from satellite-linked time depth recorders to model the proportion of time the tags were dry as a function of day-of-year, solar hour, and weather covariates (London et al. 2022). Weather included precipitation, atmospheric pressure, air temperature, and wind (Table 1 of paper). In particular, we used generalized linear mixed pseudo-models (GLMPMs; Ver Hoef et al. 2010) to model variation in haul-out behavior as a function of covariates, temporally autocorrelated random effects, and individual random effects representing heterogeneity in individual behavior. We used the 'glmmLDTS' package (Ver Hoef *et al.* 2010) to implement GLMPMs in the R programming environment (R Development Core Team 2017). We used the following covariates in GLMPMs: day-of-year, solar hour, temperature, wind speed, barometric pressure, precipitation, and latitude. We also used day-of-year:solar hour interactions to permit diurnal patterns to change throughout the year and latitude:day-of-year interactions because bearded seals occupy a substantial range and we were interested in possible differences in the timing of haul-out along a latitudinal gradient. As with Ver Hoef et al. (2014), we included linear, quadratic, and cubic effects of day-of-year. For solar hour, we adopted a continuous formulation based on the first three terms of a Fourier series. This formulation provides a flexible model while preserving the inherit circularity needed for time-of-day effects (i.e. hour 0 should be equal to hour 24). In particular, solar hour effects were represented as

$$H_t = \sum_{j=1}^{3} \alpha_{j1} \cos\left(\frac{\pi \mathrm{t}}{2(j+1)}\right) + \alpha_{j2} \sin\left(\frac{\pi \mathrm{t}}{2(j+1)}\right),$$

where $H_t$ gives the effect for solar hour $t$ and $\alpha_{jk}$ for $j$ = 1, 2, or 3 and $k$ = 1 or 2 are estimated parameters (regression coefficients).

After models were fitted to bearded seal telemetry records (see London et al. 2022 for code and data), we made predictions for time, location, and weather variables realized for each surveyed cell as $\tilde{\boldsymbol{\eta}} = 1/(1+\exp(-\boldsymbol{L}\hat{\boldsymbol{\beta}}_\eta))$, where $\boldsymbol{L}$ represents a design matrix for predictions and $\hat{\boldsymbol{\beta}}_\eta$ is a vector of parameters estimated with the glmmLDTS package. We used the delta method (Dorfman 1938) to calculate the variance-covariance matrix of predictions, $\hat{\boldsymbol{\Sigma}}_{\tilde{\eta}}$. We then computed expected logit-scale detection probability as the logit of the product of availability, nondisturbance, and thermal detection probability,

$$\boldsymbol{\mu}_{2p} = \log(\tilde{\boldsymbol{\eta}}\hat{d}\hat{b}_2/(1-\tilde{\boldsymbol{\eta}}\hat{d}\hat{b}_2)),$$

where once again $i$=2 for bearded seals. The associated covariance matrix, $\boldsymbol{\Sigma}_{2p}$, was calculated using well known formulae for the variance of products, and the delta method was once again used to transform variance to the logit scale. A logit-scale prior for bearded seal detection probability ($\boldsymbol{\nu}_2$) could then be written as

$$[\,\boldsymbol{\nu}_2] \equiv \text{Multivariate normal}(\boldsymbol{\mu}_{2p}, \boldsymbol{\Sigma}_{2p}). \qquad \text{Eq. S2}$$

#### 1.3. JOINT MARGINAL PSEUDO-LIKELIHOOD

Using bracket notation to define probability distributions (e.g. [A] gives the distribution of A and [A | B] denotes the distribution of A given B), we first wrote the joint marginal pseudo-likelihood (JMPL) for abundance and related parameters given count data as

$$[\boldsymbol{N}, \boldsymbol{\alpha}, \boldsymbol{\delta} \,|\boldsymbol{C}, \boldsymbol{\theta}] \propto \int_{\kappa}\int_{\beta}\int_{\nu}[\boldsymbol{C}|\boldsymbol{N}, \boldsymbol{p}, \boldsymbol{\theta}, \boldsymbol{\beta}][\boldsymbol{p}|\boldsymbol{\nu}, \boldsymbol{\kappa}, \boldsymbol{\theta}][\boldsymbol{\beta}|\boldsymbol{\theta}][\boldsymbol{\kappa}|\boldsymbol{\theta}][\boldsymbol{\nu}|\boldsymbol{\theta}]\Lambda(\boldsymbol{\alpha}|\boldsymbol{\theta})\mathrm{d}\boldsymbol{\kappa}\mathrm{d}\boldsymbol{\beta}\mathrm{d}\boldsymbol{\nu} \qquad \text{Eq. S3}$$

where $\boldsymbol{\theta} = \{\boldsymbol{\mu}_{1p}, \boldsymbol{\mu}_{2p}, \boldsymbol{\Sigma}_{1p}, \boldsymbol{\Sigma}_{2p}, \boldsymbol{w}_s, \boldsymbol{X}, \boldsymbol{S}, \boldsymbol{m}, \boldsymbol{A}, \boldsymbol{Y}, \boldsymbol{\delta}\}$ are the "data" input into the model (including prior means and covariance matrices), $[\boldsymbol{C}|\boldsymbol{N}, \boldsymbol{p}, \boldsymbol{\theta}, \boldsymbol{\beta}]$ is the joint probability mass function for count data, $[\boldsymbol{p}|\boldsymbol{\nu}, \boldsymbol{\kappa}, \boldsymbol{\theta}]$ is the model for detection probability (a function of both estimated and latent parameters), $[\boldsymbol{\beta}|\boldsymbol{\theta}]$ is the prior distribution for regression coefficients, $[\boldsymbol{\kappa}|\boldsymbol{\theta}]$ is the spline prior for ringed seal basking proportions, $[\boldsymbol{\nu}|\boldsymbol{\theta}]$ is a prior distribution for detection probability (including haul-out probability), and $\Lambda(\boldsymbol{\alpha}|\boldsymbol{\theta})$ is a penalty on ringed seal availability that forces it to 1.0 once the MDSDA algorithm suggests that snow melt has occurred (i.e. $\delta_{st} \geq 0$). The parameter sets $\boldsymbol{\nu}$, $\boldsymbol{\kappa}$, and $\boldsymbol{\beta}$ are integrated out of the likelihood via Laplace approximation as implemented in Template Model Builder (TMB), and can be interpreted as random effects. We term this a "pseudo-likelihood" because of the nonstandard penalty that we employed to force all ringed seals to have availability equal to haul-out proportions once $\delta_{st} > 0$:

$$\Lambda(\boldsymbol{\alpha}|\boldsymbol{\theta}) = w \sum_{k=0}^{45}(1.0 - [1 + \exp{(-f(k)]^{-1}})^2. \qquad \text{Eq. S4}$$

Here, $k$ gives different values for δ (up to the maximum observed value in aerial surveys), and $f(k)$ represents the spline formulation $\mathbf{y}\boldsymbol{\kappa}$ evaluated at its associated δ value.

Unfortunately, inference with Eq. S3 did not exhibit satisfactory performance. In particular, the prior mode for detection ($\boldsymbol{\nu}$) differed substantially from the empirical Bayes estimator of $\boldsymbol{\nu}$ associated with the S3 fitted model (see, e.g., Osgood-Zimmerman and Wakefield 2022). Because there is presumably little information in the count data alone to inform detection, this difference was likely a result of confounding between detection and habitat models (see e.g. Bravington et al. 2018). We thus developed a JMPL where detection parameters ($\boldsymbol{\nu}$) were fixed, and used it to estimate abundance and related parameters:

$$[\boldsymbol{N}, \boldsymbol{\alpha}, \boldsymbol{\delta} \,|\boldsymbol{C}, \boldsymbol{\theta}, \boldsymbol{\nu}] \propto \int_{\kappa}\int_{\beta}[\boldsymbol{C}|\boldsymbol{N}, \boldsymbol{p}, \boldsymbol{\theta}, \boldsymbol{\beta}]\,[\boldsymbol{p}|\boldsymbol{\nu}, \boldsymbol{\kappa}, \boldsymbol{\theta}][\boldsymbol{\beta}|\boldsymbol{\theta}][\boldsymbol{\kappa}|\boldsymbol{\theta}]\Lambda(\boldsymbol{\alpha}|\boldsymbol{\theta})\mathrm{d}\boldsymbol{\kappa}\mathrm{d}\boldsymbol{\beta}. \qquad \text{Eq. S5}$$

We tried different values for the weight, $w$, finding that $w = 500$ allowed the proportion of hauled out seals that are basking (i.e. not hidden lairs) to increase to an asymptote near 1.0 by the time δ = 0, while still allowing pseudo-likelihood convergence. This value was retained for all phases of estimation.

#### 1.4. Parametric bootstrap procedure

To propagate uncertainty associated with detection probability into final abundance estimates, we developed a parametric bootstrap procedure. The basic idea was to compute the variance of abundance using the well-known law of total variance:

$Var(\widehat{N}) = E(Var(\widehat{N}|\boldsymbol{\nu})) + Var(E(\widehat{N}|\boldsymbol{\nu}))$.

For each of $i = 1,2,\cdots,500$ bootstrap replicates, we simulated a sample of $\boldsymbol{\nu}_{(i)}$ from its prior distributions (i.e. Eqs. S1 and S2) and conducted joint likelihood optimization with $\boldsymbol{\nu}_{(i)}$ parameters entered as fixed values. For each bootstrap replicate, we compiled $\widehat{N}_{(i)}$, the estimate of abundance for the ith bootstrap replicate, and $Var(\widehat{N}_{(i)}|\boldsymbol{\nu}_{(i)})$, the conditional estimate of variance obtained using the inverse of the Hessian estimated during joint likelihood optimization. An estimate of variance was then constructed as

$$Var(\widehat{N}) = \overline{Var(\widehat{N}_{(i)}|\boldsymbol{\nu}_{(i)})} + Var(\widehat{N}_{(i)}). \qquad \text{Eq. S6}$$

A short simulation study (P. Conn, unpublished data) suggested that this estimator had reasonable confidence interval coverage. We used Eq. S6 to construct 95% log-based confidence intervals for total abundance (Burnham et al. 1987, Buckland et al. 2005).

#### 1.5. GOODNESS-OF-FIT

We used randomized quantile residuals (RQRs; Dunn and Smyth 1996) to assess how well our preferred model fit count data. This form of residual employs randomized 'jittering' and a transformation based on the cumulative distribution function to transform residuals from discrete space on [0,∞) to continuous space on (0,1), allowing an easier visual assessment of the fit of a model to data. We conducted separate tests for each observation type (ringed seal, bearded seal, and unknown species) and country (U.S. and Russia). For each generic count $y_k$, RQRs were generated as

$$RQR_k = F(y_k|\hat{\mu}_k, \hat{\phi}, \hat{\rho}) + u_k f(y_k|\hat{\mu}_k, \hat{\phi}, \hat{\rho}),$$

where $\hat{\mu}_k$ gives a predicted count, $\hat{\phi}$ is an estimated dispersion parameter for the Tweedie distribution, $\hat{\rho}$ is the estimated power parameter of the Tweedie distribution, and $u_k$ is a uniform random deviate. The function $F$ gives the cumulative mass function of the Tweedie distribution, while $f$ references the probability mass function associated with the Tweedie distribution.

According to this scheme, RQRs should be uniformly distributed on (0,1) if a model fits the data well. We assessed fit visually (Fig. S1) and by conducting $\chi^2$ goodness-of-fit tests. For each count and survey type, we split RQRs into 10 bins of equal length in order to compute observed and expected values needed for calculation of $\chi^2$ tests. Using this approach, p-values were 0.11, 0.14, and 0.72 for ringed seals, bearded seals, and unknown seal species in U.S. surveys, and 0.50, 0.61, and 0.83 for ringed seals, bearded seals, and unknown seal species in Russian surveys. Although none of these p-values is statistically significant at the $\alpha$=0.05 level, there is some slight indication of underfitting very large counts, particularly for U.S. observations of ringed seals (Fig. S1.1)

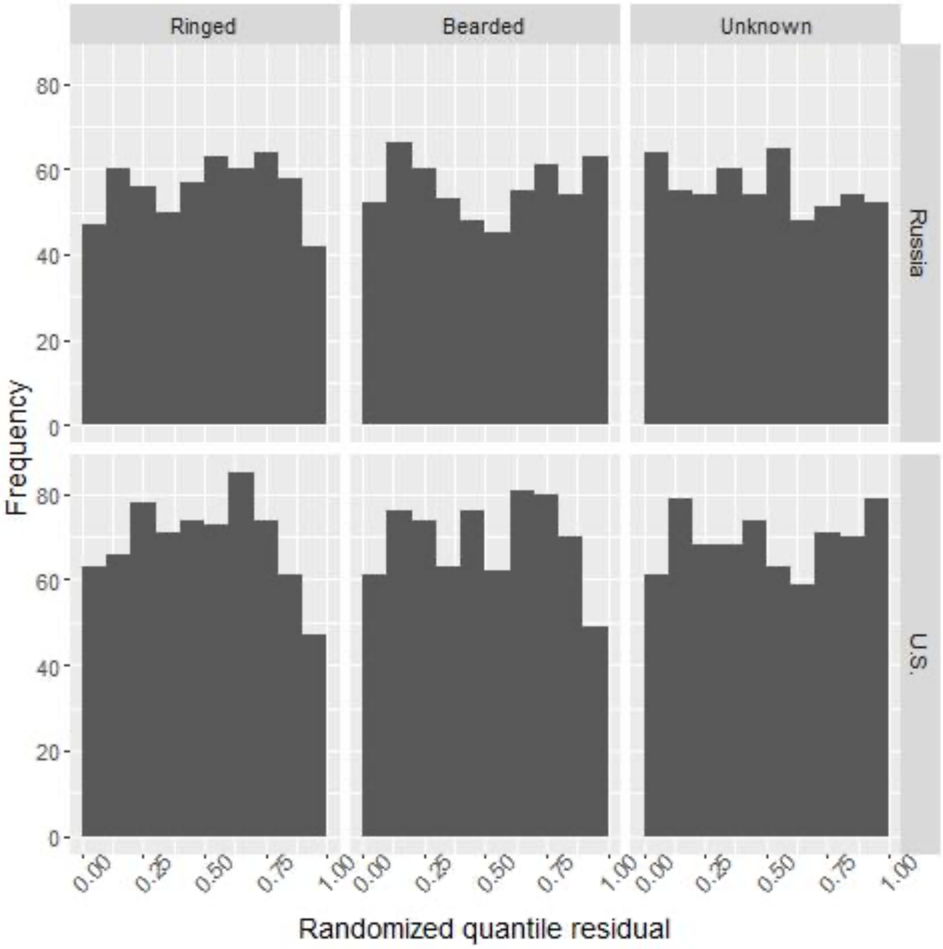


**Figure S1.** Binned randomized quantile residuals examining the fit of our preferred model to our count data, assuming counts were Tweedie distributed. For a well-fitting model, RQRs should be approximately uniformly distributed on (0,1).

### 2. Ringed seal availability modeling

In this supplement, we provide details on how we modeled ringed seal availability, $a_{st}$, the proportion of seals that are visible on ice when surveys are conducted in location $s$ at time $t$. Our basic approach is to partition availability as $a_{st} = h_{st}\alpha_{st}$, where $h_{st}$ is the proportion of seals that are hauled out, and $\alpha_{st}$ indicates the proportion of hauled out animals that are visible (i.e. not in snow lairs). We use data from ringed seals fitted with satellite-linked time-depth recorders (TDRs) to directly estimate haul-out proportions (i.e. $h_{st}$) as a function of environmental covariates. Then, covariates measured in location $s$ at time $t$ can be used to predict $h_{st}$ for times and locations surveyed during transect flights. However, these data do not provide direct information on $\alpha_{st}$, as seals can be hauled out but obscured from view when they are using subnivean lairs (Kelly *et al.*, 2006). Before describing our analysis of TDR records in further detail, we first provide more details about the conceptual problem.

#### 2.1 Conceptual model

The relationship between the proportion of ringed seals hauling out (defined as a seal being out of water) and the proportion of seals basking (defined as the proportion that are out of water *and* visible to aircraft) is summarized in Fig S2.1. The problem is that data from TDRs can be used to estimate the proportion of seals hauled out (the dashed line in Fig. S2.1), but we really need the proportion that are basking (solid line) for an aerial survey correction factor. However, we note that the aerial survey data themselves contain information about the shape of the solid line, but not the absolute value. For instance, assuming that spatially-explicit densities of ringed seals remains roughly constant over time, increases in ringed seal counts as the survey season progresses are indicative of increasing proportions of seals basking on ice (Lindsay *et al.*, In review). Our approach to modeling ringed seal detection relies on both pieces of data and the following procedure: (1) estimate the absolute value of the proportion of seals basking, using TDRs at later dates (i.e. (iii) in Fig. S2.1), and (2) estimate the shape of the availability curve (solid line in Fig. S2.1) within a larger model for seal counts. The remainder of this appendix concerns itself with a haul-out analysis for estimating $h_{st}$ from TDRs, as well as an examination of diel behavior from TDRs to inform when seals start to exhibit peak basking behavior.

#### 2.2 Analysis of ringed seal TDR data

We obtained TDR records of ringed seals from several organizations, including the U.S. Alaska North Slope Borough (NSB), the Alaska Department of Fish and Game (ADF&G), the U.S. National Oceanic and Atmospheric Administration (NOAA), and the Department of Fisheries and Oceans Canada (DFO). Capture protocols, tag descriptions, and analyses with subsets of these data are provided in Von Duyke *et al.* (2020) for NSB; in Crawford *et al.* (2012) for ADF&G; and in Harwood *et al.* (2007) for DFO.

We restricted analysis to records that were available between April and June, which is the period when seals engage in critical life history functions such as pupping, breeding, and molting. In total, records spanned the period 2005-2019 and included 54,654 hourly records from deployments on 65 individual seals. These seals provided data from the Bering, Chukchi, and Beaufort Seas (Fig. S2.2). In general, there

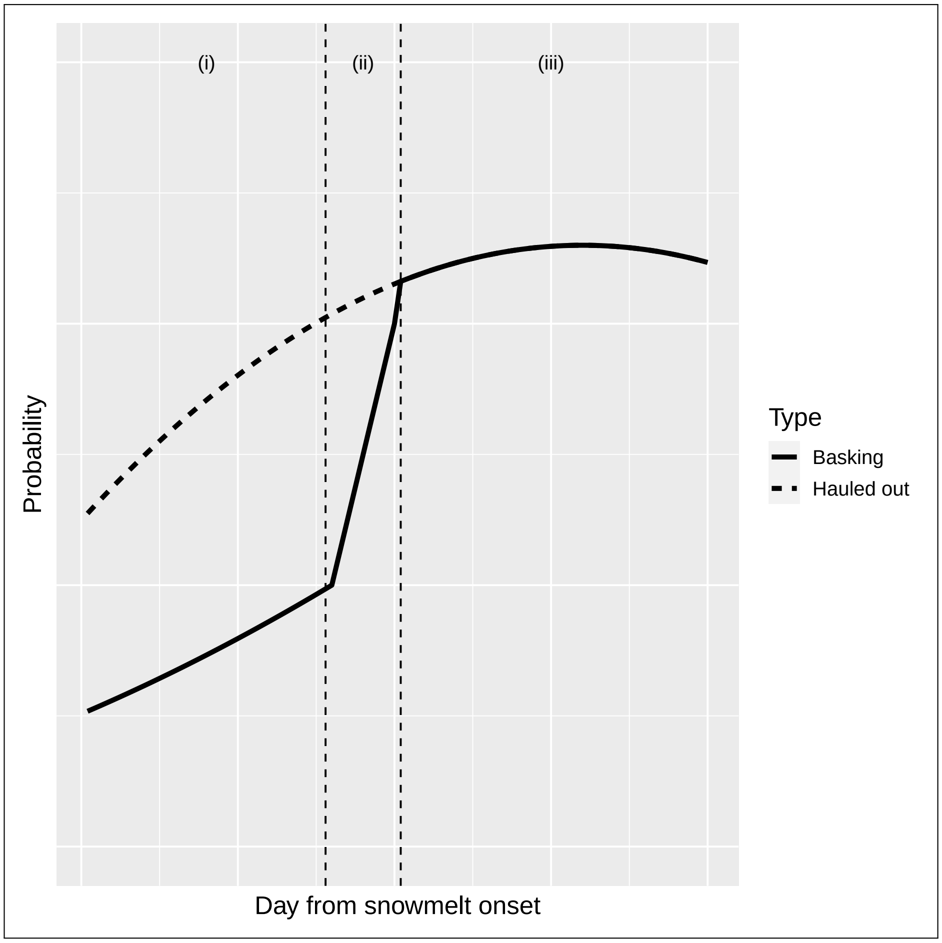


Figure S2.1: Conceptual relationship between the probability of a ringed seal basking on ice (solid line) vs the probability of ringed seals hauling out (heavy dashed line), informed by radio telemetry and direct observations of seals in the Beaufort Sea (Kelly *et al.*, 2006). Prior to transition from subnivean lairs to basking (i), many ringed seals that are "hauled out" are still in subnivean lairs and unavailable for detection. As weather warms (ii), seals start to bask on ice with higher probability, but a decreasing fraction are still hidden in lairs. Finally, once moisture saturates the snow and leads to collapse, seals enter peak basking season (iii), when hauling out is synonymous with basking.

were more records in April and early May from the Bering and Beaufort Seas and more records in late May and June for the Chukchi Sea (Fig. S2.3). We analyzed these records in several ways. First, we used generalized additive models (GAMs) to investigate the proportion of ringed seals hauled out (i.e. out of water) as a function of day-of-year and environmental covariates. Second, we used them to examine changes in diel patterns that are expected when seals transition from hauling out primarily at night when using subnivean lairs to hauling out primarily during the day to bask (Kelly *et al.*, 2006).

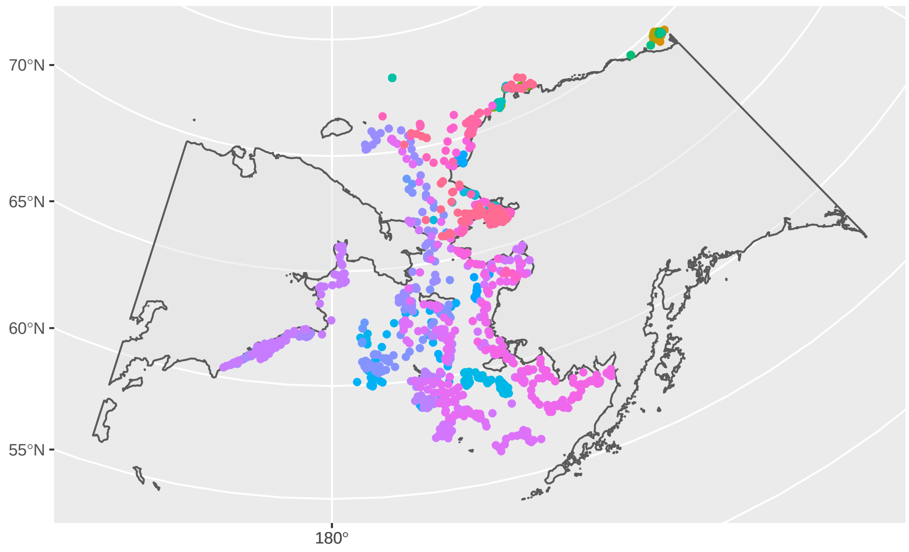


Figure S2.2: Spatial location of ringed seal haul-out records. Different colored dots represent approximate locations of individual seals.

##### 2.2.1 Haul-out analysis

To estimate the proportion of seals that were hauled out during typical survey hours ($h_{st}$), we first calculated the proportion of time each TDR was dry (all TDRs were

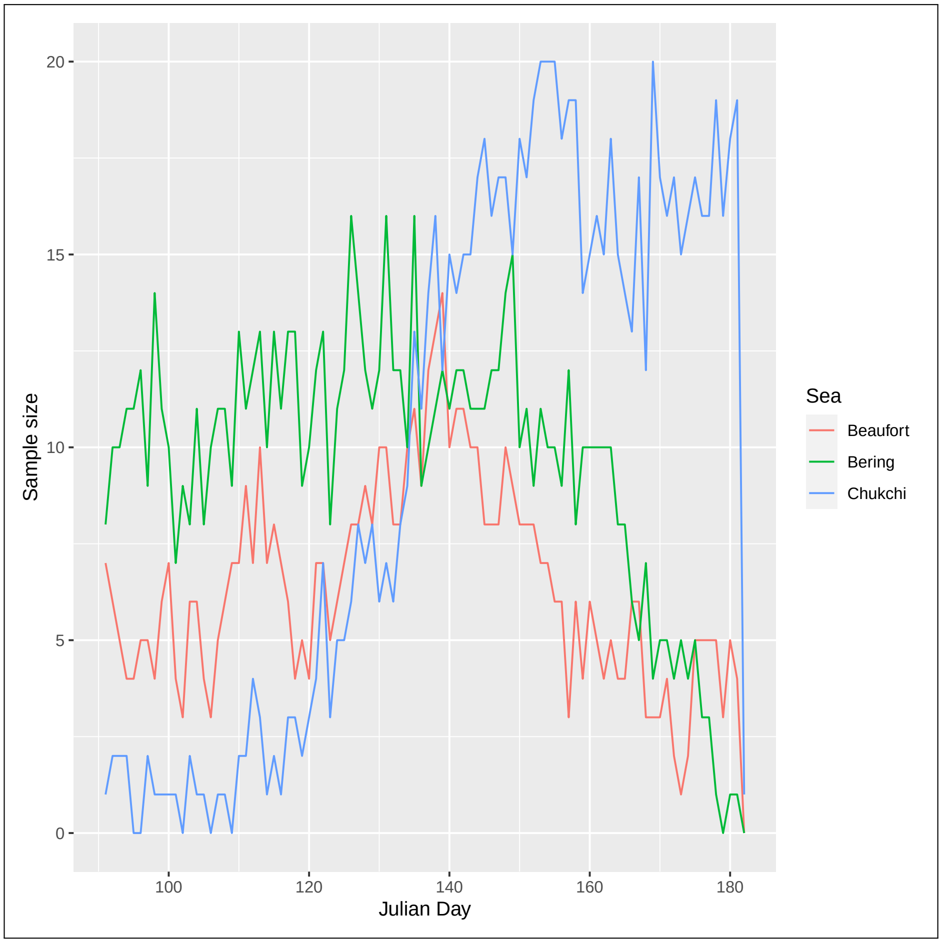


Figure S2.3: Number of ringed seals contributing haul-out records from April-June as a function of sea. For purposes of this analysis, we considered the border between the Bering and Chukchi Seas to be 65.7° N, and the border between the Chukchi and Beaufort Seas to be 156.5° W.

equipped with wet/dry sensors) for each hour between 9:00 AM and 3:00 PM local solar time as a function of Julian day and sea (Beaufort, Bering, or Chukchi). This time window corresponds to the hours where we focused Chukchi Sea aerial survey efforts in 2016 (aerial surveys in the Bering Sea in 2012 and 2013 used a similar protocol; Conn *et al.* (2014)). Restricting records to these hours, sample size is reduced to 13,670 hourly records (all 65 telemetered seals were represented). Each proportion was rounded to zero or one for use as a binomial response (87% of observations had proportions $\leq$ 10% or $\geq$ 90%). We then fit generalized additive models (GAMs; Wood, 2017) to these data using the `mgcv` package (Wood, 2017) in the R Statistical Programming Environment (R Development Core Team, 2017).

We fit several alternative GAMs to ringed seal haul-out records to examine how the timing of spring haul-out patterns varied as a function of environmental covariates. In particular, we examined whether haul-out proportions varied as a function of (1) Julian day, (2) average April-May temperature, and (3) day from estimated spring snowmelt onset. To attach temperature and snowmelt onset to individual haul-out records, we used the following procedures:

1. Predicted temperature values (at 2 m above the earth surface) were downloaded from the North American Regional Reanalysis (NARR) model produced by the National Centers for Environmental Prediction (Mesinger *et al.*, 2006) in raster format. For each year, we averaged all April and May temperature values reported between 9:00 AM and 3:00 PM local solar time for each raster grid cell (temperature values thus varied spatially and by year). Local solar time was computed using the `solaR` package (Perpiñán, 2012) in R. Hourly haul-out records were then assigned temperature values from the closest grid cell to the seal's estimated position.

2. We considered two types of snowmelt onset data derived from microwave sensors on satellites: the MDSDA algorithm (Belchansky *et al.*, 2004) and that produced by an algorithm described by Markus *et al.* (2009). In an intensive 5-yr field study in the Beaufort Sea, Kelly *et al.* (2006) demonstrated that the MDSDA algorithm was an accurate predictor of the dates of final lair use (when all ringed seals could be expected to have changed to a basking state). In particular, estimated MDSDA snow melt dates were accurate to within three days for each year with an estimated $R^2$ of 0.98. However, despite the high correlation, it is worth noting that it was based on a small sample ($n = 5$) and that multiple correlations were examined (the MDSDA-final date of emergence correlation being the highest). We suspect that this correlation would be lower had more years data been available. Despite these considerations, it remained the best candidate available - it's only trouble being that it must be calculated directly using raw satellite microwave sensor data. The snow melt product using the Markus *et al.* (2009) algorithm, on the other hand, is readily accessible and we downloaded it directly from the National Snow & Ice Data Center (Steele *et al.*, 2019). For each year, these two algorithms produced spatially explicit (raster) estimates of the Julian day of snow melt onset on a 25 $\times$ 25 km resolution. However, melt onset values were often missing in locations close to land; we interpolated these values using geostatistical kriging as employed by the `automap` package in R (Hiemstra *et al.*, 2008). For each hourly haul-out record, we attached the closest melt product value (spatially) to the estimated animal position. We then calculated the day from melt onset as

$$\delta_{i,t} = jday_{i,t} - melt_{s(i,t),y(i,t)}, \quad (1)$$

where $jday_{i,t}$ is the Julian day associated animal $i$ at time $t$, and $melt_{s,y}$ gives the Julian day of melt onset in location $s$ in year $y$ (which are themselves a function of the position of seal $i$ at time $t$).

Modeling was conducted in several stages. First, because we were concerned that temporal autocorrelation in responses would lead to over-dispersion relative to binomial variance, we estimated a variance inflation factor, $\phi$, using the "quasibinomial" family option available in mgcv. We did this for our two most highly parameterized models (see below), obtaining estimates of $\phi = 3.6$ in both cases. Second, we conducted model selection using qAIC, a version of Akaike's information criterion (Akaike, 1973) adapted for overdispersion (Burnham & Anderson, 2002), to assess the relative ability of the two remotely sensed spring snowmelt onset products to predict haul-out probabilities. In this stage, we fitted two GAMs, each of which included (i) either of the two snow melt products, and (ii) Julian day and average April-May temperature as additional predictor variables. Initial data exploration indicated qualitatively different trends in haul-out distributions between seals in the Bering Sea from those in the Chukchi/Beaufort Seas, so we estimated separate effects for each of these areas (here defined as south or north of the Bering Strait, respectively). Finally, in the third stage of modeling, we sequentially dropped individual terms from the highest-ranked snow melt product model to examine their effect on predictive performance (similar to backwards selection). For all stages, we used a thin plate spline basis (with default degrees of freedom) in `mgcv` to model smooth effects of covariates on haul-out proportions.

| Model | LogL | k | qAIC | $R^2$-adj |
|---|---|---|---|---|
| Markus*sea + jday*sea + temp | -5298.60 | 26.57 | 3003.01 | 0.39 |
| Markus*sea + jday*sea + temp*sea | -5299.67 | 26.75 | 3003.97 | 0.39 |
| Markus + jday*sea + temp*sea | -5308.08 | 26.90 | 3008.95 | 0.39 |
| Markus*sea + jday + temp*sea | -5314.52 | 26.78 | 3012.29 | 0.39 |
| MDSDA*sea + jday*sea + temp*sea | -5318.83 | 27.39 | 3015.92 | 0.39 |
| Markus*sea + jday*sea | -5364.61 | 18.41 | 3023.46 | 0.38 |
| Markus*sea + temp*sea | -5433.25 | 19.03 | 3062.91 | 0.37 |
| jday*sea + temp*sea | -5440.20 | 18.79 | 3066.28 | 0.37 |

Table 1: Models fit to ringed seal haul-out data, together with fitted log-likelihood values ("LogL"), estimated number of parameters ($k$), qAIC score, and adjusted $R^2$ value ($R^2$-adj; as output by `mgcv`). A lower qAIC score indicates a model with better predictive performance (models within 2.0 qAIC points of each other are often interpreted as having equal support). Note that the `mgcv` R package includes penalization on smooth terms, which induces a non-integer parameter count.

Comparing GAM fits by qAIC suggested better predictive performance of models that included Julian day, spring temperature, snow melt onset data generated with the Markus *et al.* (2009) algorithm, with melt and Julian day effects varying by sea (Bering vs. Chukchi/Beaufort) (Table 1). There was approximately equal support between models with and without an interaction between spring temperature and sea. However, even though qAIC scores differed substantially, this translated into only minor differences in $R^2$. Nevertheless, we proceeded with the highest ranked AIC model to generate predicted haul-out values as a function of Julian day, temperature, and snow melt onset values.

The highest ranked GAM model predicted that ringed seals gradually increased the proportion of time hauled out during typical survey hours as Arctic spring progressed (Figs. S2.4-S2.5). Under average environmental conditions, peak haul-out proportions in the Chukchi and Beaufort seas were predicted to reach 0.8-0.9 in late May and early June, and to peak earlier and at much smaller values in the Bering Sea. There was considerable year-to-year variation in the timing and magnitude of predicted haul-out proportions. This variation has potentially large ramifications for correction factors applied to aerial surveys, including for 2016 surveys in the Chukchi Sea.

Peak haul-out proportions estimated here are similar to, but slightly larger than, peak basking proportions ($\approx 65\%$) used in a previous analysis of Chukchi Sea ringed seal aerial surveys (Bengtson *et al.*, 2005) based on a smaller sample of telemetered seals. Our estimates are also similar to the value of 0.75 obtained during a 5-yr, intensive observation and radio-telemetry study in the Beaufort Sea (Kelly *et al.*, 2006).

Although we have not presented data on the age of tagged animals (owing to missing records and a variety of ways of summarizing the size, age, or age class of individuals), it was clear from inspecting data that tagged seals in the Bering Sea were younger than those in the Chukchi or Beaufort Seas. Relatively low predicted haul-out proportions in this region may be because the seals represented in this sample were comparatively free of pupping and breeding obligations. However, an important question (and one that we cannot answer with the data at hand) is whether this sample is representative of the population of ringed seals in the Bering Sea in April and May. Crawford *et al.* (2012) suggested that seals in the Bering Sea on the whole may be biased towards younger age classes. Alternatively, Kelly (2022) argued that this finding could be an artifact of the tagging process, with opportunistic summer sampling of ringed seals near Kotzebue, Alaska disproportionately sampling younger, transient seals that are more likely to move to the Bering Sea. The implications for correction factors in the Bering Sea is quite profound; for instance if predictions from Chukchi-Beaufort seals were used as correction factors for the Bering Sea in (instead of our tagged sample of subadult-biased seals that move to the Bering Sea), abundance estimates would likely be underestimated by a factor of $2-3$ in May.

##### 2.2.2 Diel behavior and den emergence

As a second way of viewing our data, we examined the diel behavior of seals as a function of MDSDA snow melt onset date using rose plots (Fig. S2.6). Previous research in the Bering, Chukchi, and Beaufort Seas (Kelly *et al.*, 2006; Crawford *et al.*, 2019; Von Duyke *et al.*, 2020) indicated that seals using subnivean lairs primarily hauled out at night, switching to hauling out during the day when emerging from lairs. We observed a similar pattern for telemetered ringed seals in our study (Fig. S2.6), which used many of the same haul-out records. Importantly, Kelly *et al.* (2006) found that the MDSDA algorithm for snow melt onset was an accurate predictor for *final* lair use (meaning the day the last seal in their study emerged and changed to the basking state). Reflecting on our conceptual model, it thus seems reasonable that the transition to basking behavior (i.e. from (ii) to (iii) in Fig. S2.1) is complete when $\delta_{s,t} = 0$. We thus propose the following model for use in analysis of aerial survey counts:

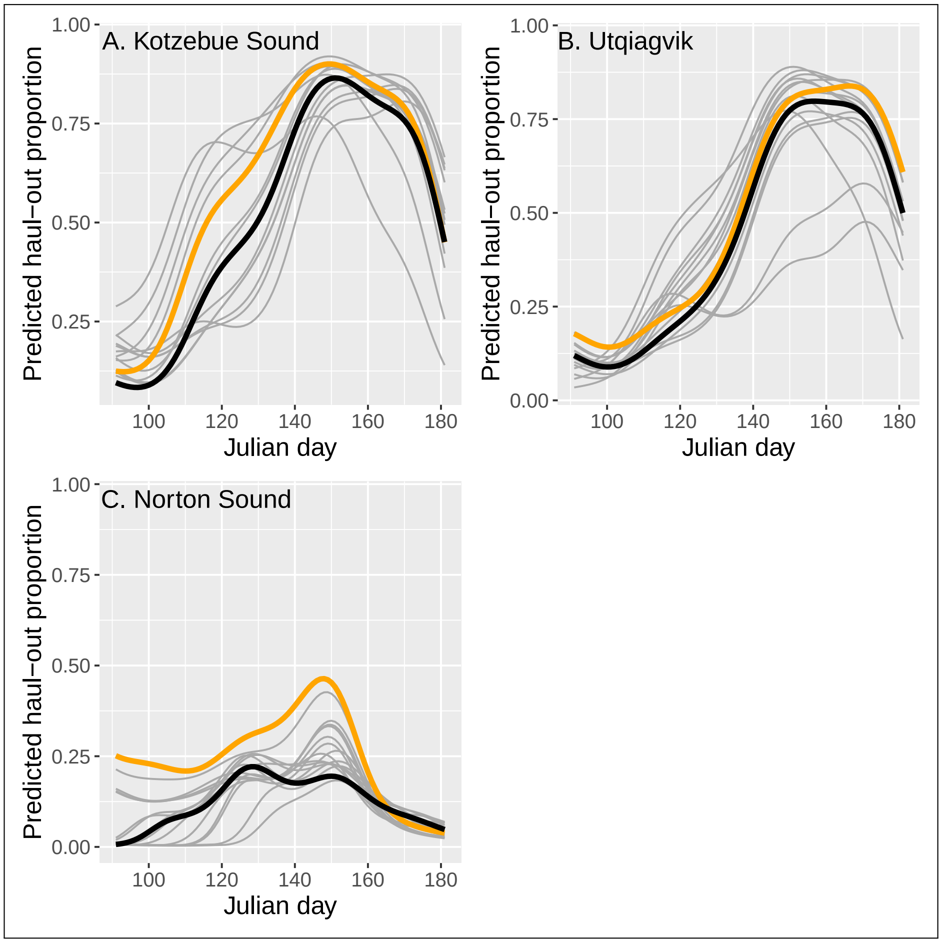


Figure S2.4: Proportion of ringed seals hauled out of water as a function of location, year, and Julian day as predicted by the highest ranked GAM model fitted to ringed seal satellite telemetry records. Light gray lines provide predictions for individual years (2005-2019), with 2016 (the year of Chukchi Sea aerial surveys) in gold, and the overall mean prediction (taken across years) in black. Separate predictions were made for three representative areas with different snow melt onset and temperature values: Kotzebue Sound, Alaska (66.52° N, 162.75° W); Norton Sound, Alaska (64.5° N, 162.75° W); and near Utqiagvik, Alaska (71.31° N, 156.71° W).

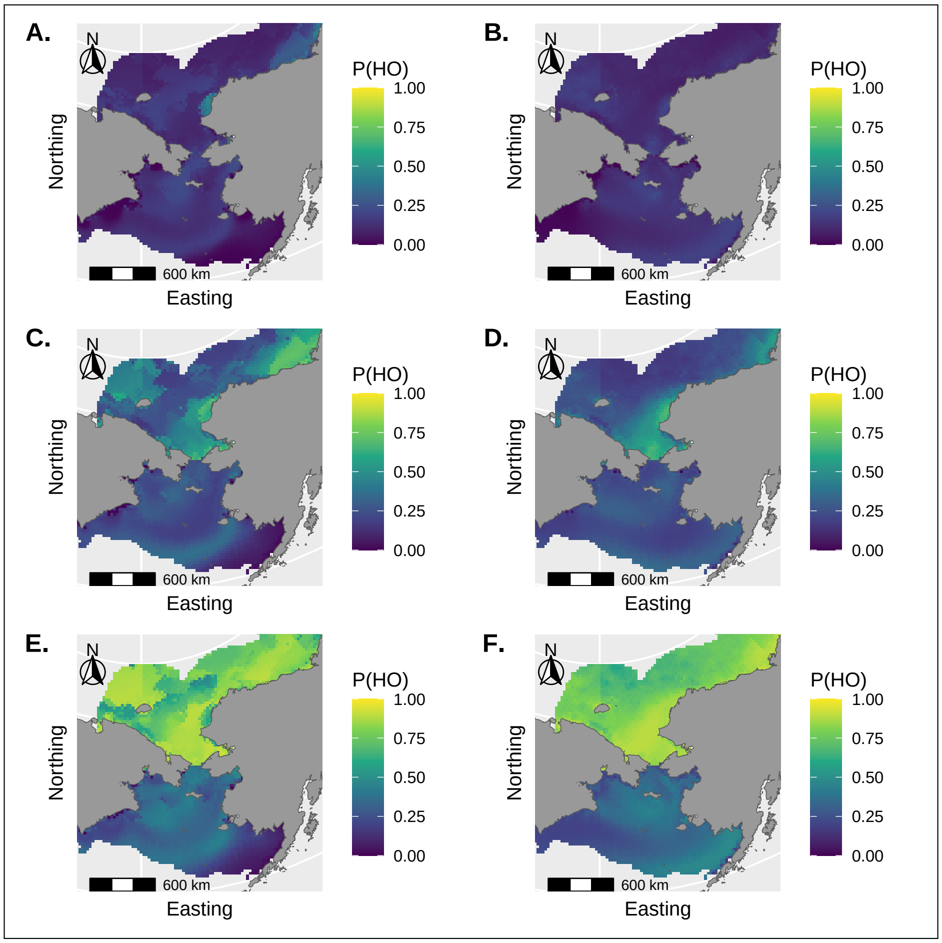


Figure S2.5: Proportion of ringed seals predicted to haul-out as a function of environmental conditions on specific Julian days. Maps on the left (A, C, and E) give predictions for 2016, while maps on the right (B,D,F) give predictions averaged over years (2005-2019). The first row (Plots A-B) are for Julian day 100 (April 9 for 2016); the middle row (Plots C-D) are for Julian day 125 (May 4 for 2016); the last row (Plots E-F) are for Julian day 150 (May 29 for 2016).

$$a_{s,t} = \begin{cases} h_{s,t} & \delta_{s,t} \geq 0 \\ h_{s,t}\alpha_{s,t} & \delta_{s,t} < 0 \end{cases} \tag{2}$$

To impart further structure on the proportion of hauled out animals that are visible to aircraft, we suggest writing $\alpha_{s,t}$ as an increasing function of $\delta_{s,t}$. For purposes of this paper, we use a logistic function, constrained so that $\alpha_{s,t} \rightarrow 1.0$ as $\delta_{s,t} \rightarrow 0$ (see main manuscript text).

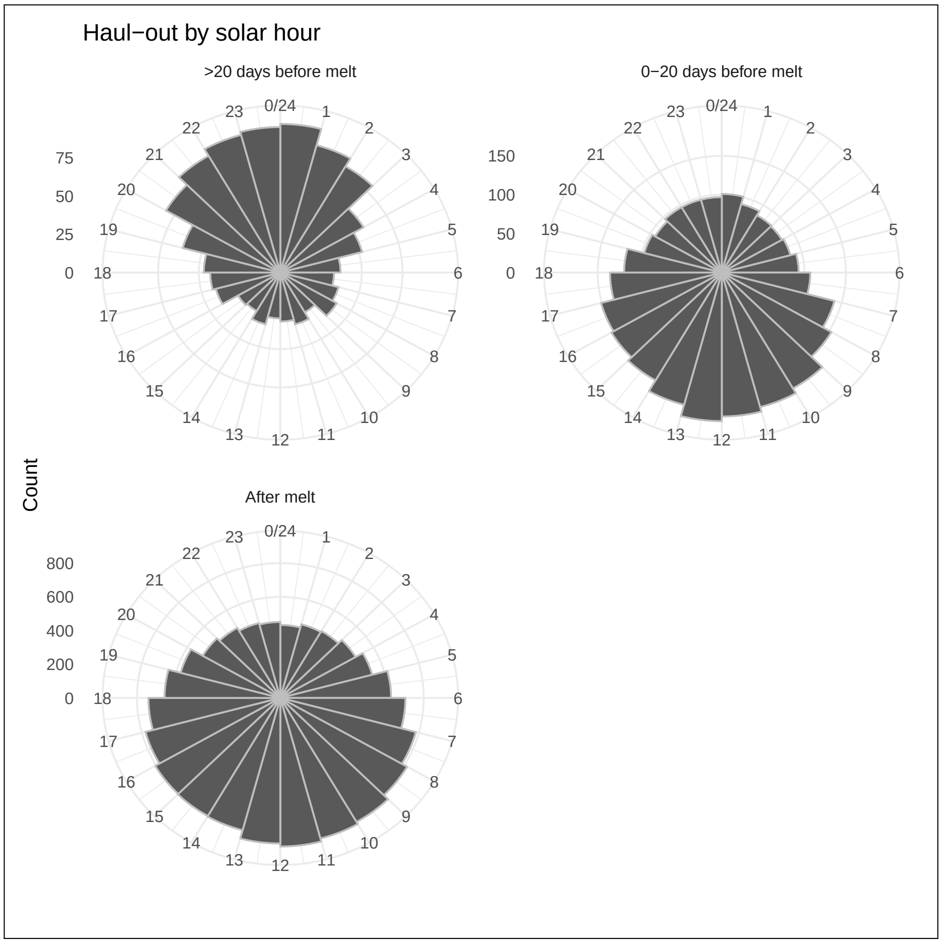


Figure S2.6: Rose plots summarizing the distribution of hour-of-day (local solar time) that telemetered ringed seals were hauled out, broken down by day from snow melt onset as calculated with the MDSDA algorithm. Twenty of more days before melt onset, seals hauled out more often at night (consistent with behavior when seals use subnivean lairs), shifting to primarily hauling out during the day as snow melt onset approaches (consistent with basking behavior).
